# Convergent gene expression highlights shared vocal motor microcircuitry in songbirds and humans

**DOI:** 10.1101/2022.07.01.498177

**Authors:** Gregory L Gedman, Matthew T. Biegler, Bettina Haase, Morgan E. Wirthlin, Olivier Fedrigo, Andreas R. Pfenning, Erich D. Jarvis

**Affiliations:** The Rockefeller University; New York, NY, 10065, USA; Howard Hughes Medical Institute; Chevy Chase, MD 20815, USA; Department of Computational Biology, Carnegie Mellon University; Pittsburgh, PA 15213

## Abstract

Vocal learning is a skilled motor behavior observed in several mammalian and avian species and is critical for human speech. While convergent gene expression patterns have highlighted similar primary motor and striatal pathways for vocal imitation in songbirds and humans, the extent of molecular and circuit convergence remains unresolved. Here we profiled the four principal song nuclei of the zebra finch (HVC, LMAN, RA, Area X) and their surrounding brain regions using RNA-Seq and compared them with specialized markers we identified for human speech brain regions. Expanding previous work, both songbird RA and HVC exhibited convergent specialized gene expression of ∼350 genes with human laryngeal sensorimotor cortex. The songbird HVC_RA_ intratelencephalic (IT) neurons were the predominant cell type that was convergent with human, specifically layer 2/3 IT neurons, while the songbird RA extratelencephalic (ET) projection neurons exhibited convergent expression with human layer 5 ET projection neurons. The molecular specializations of both songbird LMAN and human Broca’s area were more unique to each species. These findings demonstrate the extent of convergent molecular specializations in distantly related species for vocal imitation and emphasize important evolutionary constraints for this complex trait.

**One-Sentence Summary:** Our data provide hundreds of candidate genes to study the molecular basis and evolution of song and speech across species.

## Main Text

Across the animal lineage, the neural control of musculature is an essential component for survival and interaction with one’s environment. A diverse range of vertebrate cortical motor systems can be found in nature, evolving either through homologous inheritance from a common ancestor or through convergence in independent lineages (*1*). Convergent evolution of cortical motor circuits implies great functional importance and limited mechanisms for complex nervous systems to accomplish a particular motor task.

Vocal learning is one example where convergent motor brain regions can give rise to a convergent motor phenotype. Vocal learning is the ability to imitate sounds through social exposure. Vocal learning has been observed in several diverse mammalian (humans, bats, cetaceans, pinnepeds, and elephants) and avian (songbirds, parrots, and hummingbird) lineages, and is a critical component of human spoken language (*2*). Of those species studied, all have evolved a specialized forebrain sensorimotor learning circuit that is absent or very rudimentary in their closer vocal non-learning relatives (*3*). In songbirds and humans, several regions in this convergent vocal motor circuit also exhibit convergent molecular specializations, including the songbird robust nucleus of the arcopallium (RA) with the human laryngeal sensorimotor cortex (LSMC), and songbird Area X with a part of the human anterior striatum (ASt) active during speech production learning (**Fig. 1**) (*4, 5*). The premotor components of the songbird (HVC, LMAN) and human (Broca’s, SMA) vocal learning/speech circuit were either found to have weak or no significant shared gene expression specializations, despite competing hypotheses on shared connectivity and/or function (*3, 6-10*). Technical limitations could explain this lack of shared molecular specializations, including an incomplete sampling of the transcriptome using oligo microarrays, insufficient sampling of surrounding control brain regions, and use of a “winner-take-all” approach that limited more than one brain region from one species matching the same brain region in another (*4*). Alternatively, this apparent lack of convergence could simply reflect real biological differences across vocal learning species.

**Fig 1:**
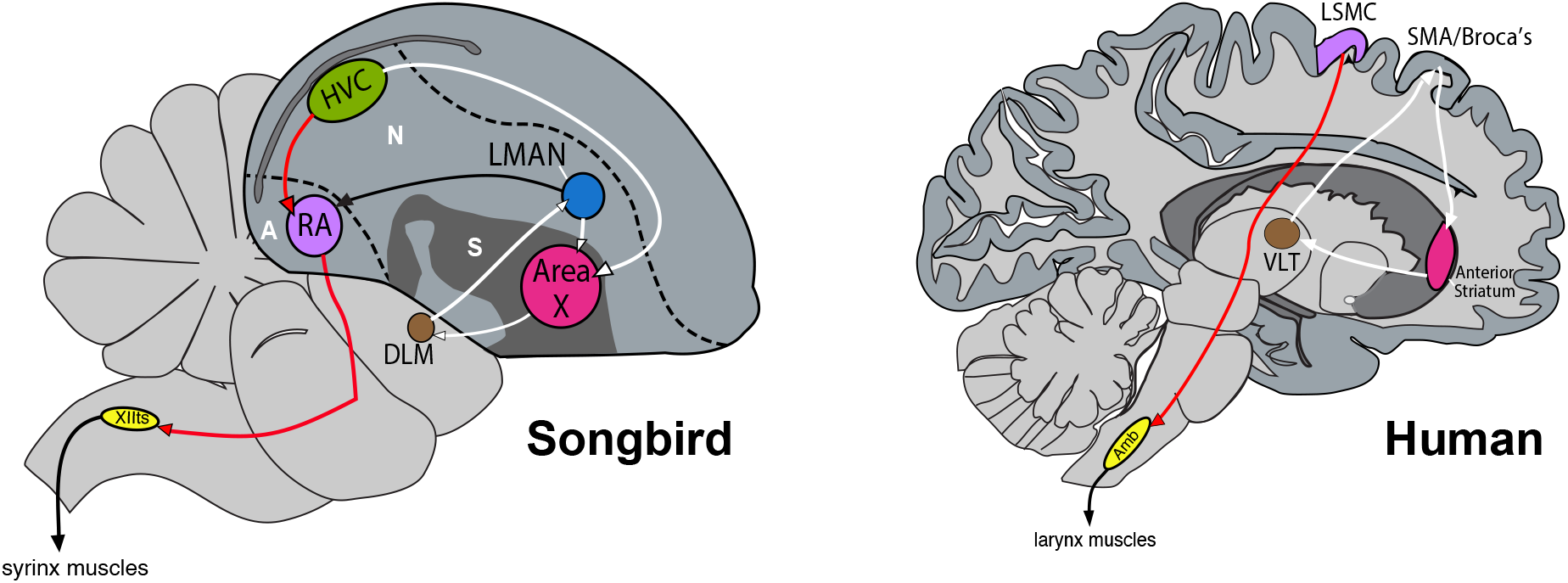
Songbird and human circuit and molecular convergence for vocal imitation. Both the species exhibit convergent neural circuitry for the control of vocal imitation, with a cortico-striatal-thalamic loop (white arrows) and a direct cortical connection to the motoneurons controlling the vocal organ (red arrows). Previous work (*4*) found that this circuit convergence is accompanied by molecular convergence between songbird RA and human LSMC (purple) as well as songbird Area X and human anterior putamen (pink). Importantly, no significant molecular analog of songbird HVC or LMAN was found in the human brain. N: Nidopallium. A: Arcopallium. S: Striatum.

Here, we set out to address these limitations and test the extent of molecular convergence in songbird and human vocal learning/speech circuitry. We profiled the whole transcriptome of songbird (zebra finch) posterior forebrain pathway (PFP) nuclei HVC and RA, anterior forebrain pathway (AFP) nuclei LMAN and Area X, as well as their adjacent non-vocal motor brain regions, using bulk RNA sequencing (RNA-Seq). We compared our RNA-Seq expression data with human brain microarray (*11*) and single-nuclei (*12*) RNA-Seq datasets from the Allen Institute for Brain Science, as well as single nuclei data from songbird HVC and RA (*13*), using gene set enrichment analyses allowing a one-to-many mapping of brain regions or cell types across species. We confirmed and more strongly support a convergent specialized molecular relationship between songbird RA and human LSMC regions, and songbird Area X and human ASt. We found novel molecular convergence between songbird HVC and human LSMC, but no strong relationship between songbird LMAN and human Broca’s. Both HVC and RA demonstrate molecular convergence with specific cell classes within human motor cortex, suggesting a convergent microcircuitry governing learned vocal production. We present a more holistic hypothesis of molecular convergence in motor circuits for song and speech in vocal learning birds and humans.

### The zebra finch song system is distinguished by several thousand specialized genes

We used laser capture microscopy to precisely isolate the four principal song nuclei (HVC, LMAN, RA, Area X) and their non-vocal motor surrounds (**Fig. S1**), and then profiled their transcriptomes using bulk RNA sequencing (Methods). Utilizing a newly assembled and annotated zebra finch genome (*14*), we then performed differential gene expression analysis between each of the song nuclei and their adjacent non-vocal motor brain regions, controlling for possible batch effects (**Fig. S2-4, Table S1**). Remarkably, we found that ∼25% of the annotated transcriptome (n = 5473 genes) exhibited specialized up- or down-regulation in one or more of the song nuclei relative to their adjacent non-vocal motor brain regions, ranging from ∼1000-3800 genes each (HVC = 1542; LMAN = 3878; RA = 1002; Area X = 1052; **Fig. 2A**). Despite these large differentially expressed gene sets, only 0.5% (n = 26) were similarly specialized to all nuclei, and 1% (n = 55) to all pallial/cortical nuclei, whereas most genes (∼30-60%) were uniquely specialized within one song nucleus each (**Fig. 2B**). In addition, we noted some similarity in the specialized gene expression exclusively found between the PFP (1.2%; n = 67) and AFP (3.9%; n = 216) nuclei, suggesting distinct molecular properties in each anatomical pathway. The nidopallial nuclei HVC and LMAN shared the largest overlap in molecular specializations (12.7%; n = 697), likely due to their shared nidopallial origin. To assess the accuracy of our differential gene expression analysis, we compared our specialized gene lists for each nucleus with validated markers of the song system by *in situ* hybridization data from our lab (n = 32 genes), the zebra finch expression brain atlas (ZeBrA, n = 142), and other published studies (n = 4) (*4, 15-17*). Of the genes examined across all nuclei (n = 178), we found high concordance (∼72-89% true positive rate) between our RNA-Seq data and *in-situ* hybridization gene expression patterns (**Fig. 2C, Fig. S5, Table S2)**, suggesting our RNA-Seq data are an accurate representation of the molecular specializations in the zebra finch vocal learning circuit. Together, these findings highlight a 5-to 10-fold higher degree of specialization distinguishing each song nucleus from its non-vocal motor surround than previously reported (*4*). These improvements are presumably due to profiling the entire transcriptome, including additional surrounding brain regions, and utilizing a higher-quality reference genome with more completely annotated genes.

**Fig 2:**
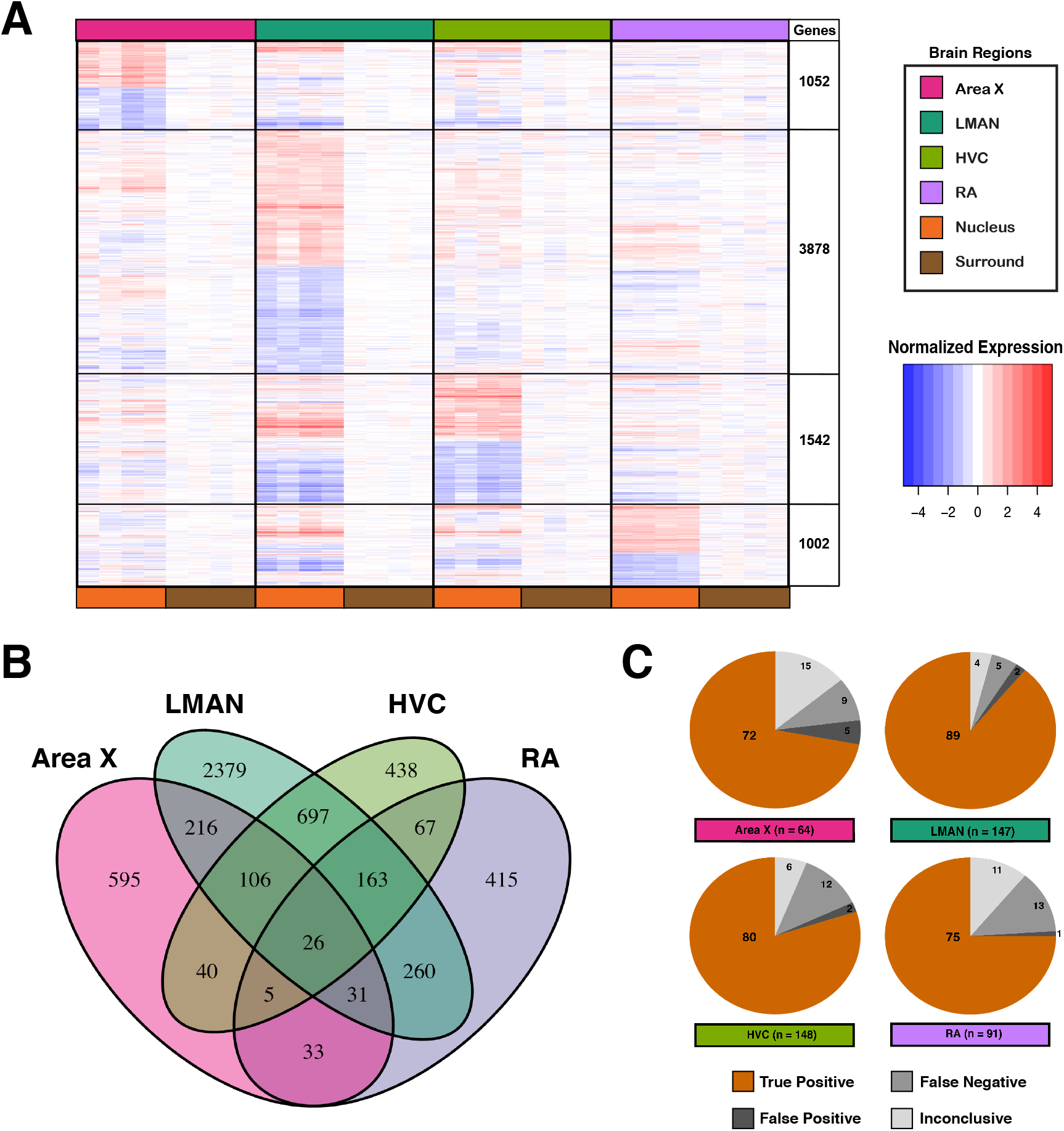
Molecular specializations of the zebra finch song system. **(A)** Heatmap of specialized genes (n=5473) between Area X, LMAN, HVC, and RA relative to their non-vocal motor surrounding regions. Each column is a replicate for the indicated brain region (color key) and each row is z-score normalized expression of a given gene. **(B)** Venn diagram quantifying overlap of molecular specializations across the song system. **(C)** Summary of in situ hybridization validations of differentially expressed genes (n = 178) in the zebra finch song system. Our gene expression data exhibit a high concordance rate with *in situ* expression patterns.

### Gene expression specialization distinguish the human speech system from the whole cortex

To test for molecular convergence across species, we first set out to define the molecular specializations for key nodes of the human spoken language network relative to the rest of the cortex as well as within the striatum (See Methods). We utilized the Allen Brain Expression Atlas, a comprehensive coordinate-based microarray dataset comprising 3681 samples from 6 postmortem brains (*11*, **Table S3**), which allowed us to select gene expression data from brain regions demonstrating activation during human speech production (*5, 9, 18-20*). These included human speech regions hypothesized to be functionally and molecularly analogous to different songbird song nuclei, namely the LSMC, SMA, and Broca’s area in the cortex, and ASt in the striatum (*3, 6-10*). Each of these broad regions consists of several subregions activated during different spoken language tasks (*5, 9, 20-23*), so additional specialized gene sets were generated using samples in MNI coordinates showing speech-related activity (**Fig. S6, Table S4)**. We found distinct gene sets with specialized expression in each human speech brain region or subregion (median = 1330, **Table S5**). Unlike in songbird, molecular specializations defining human cortical LSMC regions shared the most gene markers with subcortical specializations of ASt (**Fig. S7**). This is consistent with recent evidence of shared transcriptional networks in functionally coupled cortico-striatal brain networks, possibly driven by specific interneuron subtypes (*24*). These specialized gene sets defining key nodes in the speech motor system and their subregions were used to test for molecular convergence with songbird vocal motor nuclei.

### Robust molecular convergence between songbird and human song and speech pathways

To test for molecular convergence across species, we conducted gene set enrichment for significant overlap (convergence) in songbird and human molecular specializations using a hypergeometric test (**Fig. S8**), restricting the analysis to one-to-one orthologs in both species (n = 11,475 genes). Instances where the genes sets exhibited significant overlap (q < 0.05) greater than chance (observed (O) /expected (E) > 1) were interpreted as molecular convergence (**Fig. 3**). We confirmed the previously observed significant molecular similarity between songbird RA with human LSMC regions (**Fig. 3A,F**; n = 155 genes; q = 0.04, O/E = 1.16) and songbird Area X with human ASt subregions (**Figs. 3D, S9**; n = 118 genes; q_Cd/Pu_ = 0.015/0.023; O/E_Cd/Pu_ = 1.34/1.24). These relationships serve to validate our approach as well as triple previous numbers of convergently expressed genes (**Table S6**). Interestingly, we discovered that songbird HVC also shares robust molecular specialization with the human LSMC regions (**Figs. 3A**; q = 4.66e-05, O/E = 1.49), but with a largely distinct gene set from RA (**Fig. 3F**). Such robust molecular similarity found between HVC/RA and LSMC was not found in the adjacent human non-vocal sensorimotor (torso) cortex regions (**Fig. 3A**). In contrast, while HVC and LMAN exhibited a weaker signal of molecular convergence to SMA (**Fig. S10**), as well as HVC to Broca’s area (**Fig. S11**), both premotor nuclei also shared molecular similarity to non-vocal premotor cortex, suggesting the observed similarity is not specific to vocal learning/speech brain regions (**Fig. 3C;** q = 2.9e-03; O/E = 2.25). These data demonstrate that songbird HVC and RA share the greatest gene expression analogy to human LSMC, while LMAN shares expression similarities to the premotor cortex in general.

**Fig 3:**
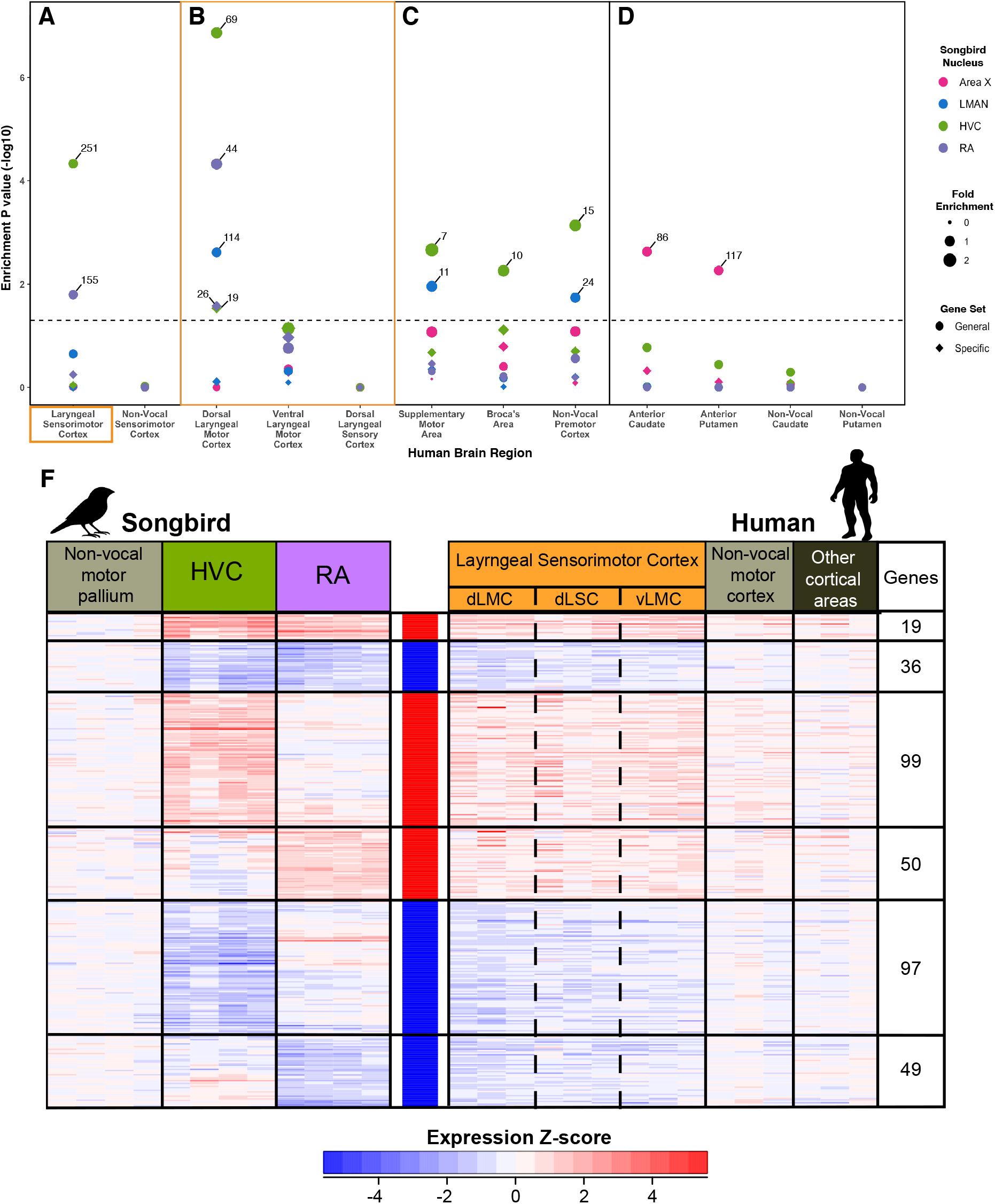
Molecular convergent between zebra finch HVC/RA and human LSMC. (**A-D**) Results of hypergeometric test for significant overlap in specialized gene expression. Gene sets defining each human brain region were independently tested against molecular specializations of each zebra finch song nucleus. Significant overlaps are those that pass FDR < 0.05 (dotted line). Size of the circle is the fold enrichment of the observed overlap over expected chance. Specialized gene sets from songbird Area X (pink), LMAN (blue), HVC (green), and RA (purple) were generally defined relative to their surrounding non-vocal motor brain regions (circles) or specifically by genes uniquely specialized to a given nucleus and no others (diamonds). Test regions include: **(A)** human laryngeal sensorimotor cortex (LSMC) and adjacent non-vocal (torso) sensorimotor cortex; (**B)** each LSMC subregion individually (dLMC, vLMC, and dLSC); **(C)** human supplementary motor area (SMA); Broca’s area, and non-vocal premotor cortex (BA6); and **(D)** human anterior caudate and putamen, as well as adjacent non-vocal motor portions of each. **(F)** Heatmap visualizing z-score normalized expression of the convergent gene set shown in **(A)** for songbird HVC/RA (left) and human LSMC (right). Each column is a replicate of the above brain region. Each row is a gene exhibiting convergent expression across songbird and human vocal motor regions. Non-vocal motor regions for each species and other cortical regions (human) are included as reference. A subset of the three subregions of the LSMC are delineated (dotted lines).

The laryngeal sensorimotor cortex (LSMC) is comprised of three anatomically separated subregions with functional distinctions (**Fig. S6)**. These include the ventral laryngeal motor cortex (vLMC), dorsal laryngeal somatosensory cortex (dLSC) and adjacent dorsal laryngeal motor cortex (dLMC;), a more functionally and evolutionarily novel region controlling human speech pitch and prosody (*18-20, 22, 25, 26*). Our prior analyses found that songbird RA exhibited greater molecular convergence with human dLMC compared to vLMC (*4*), so we hypothesized that songbird HVC might also exhibit more molecular convergence with the dLMC. To test this hypothesis, we compared the molecular specializations of each LSMC subregion with those of the zebra finch song system. We found evidence of clear regional specificity, with songbird HVC (q = 1.09e-07; O/E = 1.95), RA (q = 5.68e-05; O/E = 1.89), and LMAN (q = 2.59e-03; O/E = 0.96) exhibiting convergent molecular specializations with human dLMC (**Fig. 3B, circles**). None of these nuclei exhibited such a significant match to vLMC or dLSC alone. Yet when vLMC and dLSC are included with dLMC, additional genes show convergence with HVC and RA, indicating a weaker convergence with vLMC and dLSC. Since HVC and LMAN share a high degree (70%) of similarly specialized genes (**Fig 2B**), we hypothesized that the molecular convergence between LMAN and dLMC could be driven by shared gene sets with HVC. To test this, we examined unique gene sets for each nucleus, which contain genes exhibiting specialized expression to that nucleus and no other song nucleus. We found that the unique specialized gene sets from HVC and RA still exhibited significant molecular convergence with human dLMC, while the unique set from LMAN did not (**Fig. 3B, diamonds**; q_HVC/RA_ = 0.03; O/E = 1.49-1.56). Importantly, no songbird nucleus exhibited convergent molecular similarity to the non-vocal motor sensorimotor cortical regions, suggesting the observed convergence is specific to vocal motor pathways. Together, these data suggest that the songbird vocal motor nuclei, HVC and RA, demonstrate molecular convergence with human LSMC, specifically the more evolutionarily novel dLMC region.

### Songbird HVC and RA are represented as an analogous microcircuit in human LSMC

We next asked how two spatially and functionally distinct songbird vocal nuclei (HVC and RA), with distinct projection neuron types, can exhibit molecular similarity to the same human vocal LMC control region. One hypothesis is that these two songbird regions exhibit convergence to different cell types within the same human cortical column. Consistent with this idea, we noted that HVC and RA matched LSMC expression with largely distinct genes sets (**Fig. 3F**), suggesting that each nucleus is drawn to a specific expression pattern (i.e. cell types) within the heterogenous LSMC transcriptome.

To test this idea, we first asked if there are any specific cell types within songbird HVC and RA that strongly express genes exhibiting convergence with human LSMC. We made the comparison with genes convergent with the combined LSMC regions as opposed to dLMC alone, since the former included more significantly specialized convergent genes. Each songbird nucleus has several projection neurons types and local interneuron types (*28*). We utilized single nuclei RNA-Seq dataset of zebra finch HVC and RA (*27*), and correlated their expression levels to normalized fold changes of convergent genes shared between HVC/LSMC and RA/LSMC (Methods). We found that the HVC/LSMC convergent gene set most strongly correlated with the two types of mature HVC to RA projection neurons (HVC_RA1_, HVC_RA2_), which are known to control production of learned vocalizations, with strong anti-correlations to other HVC neuron classes (**Fig. 4A**). Similarly, the RA/LSMC convergent gene set most strongly correlated with one subclass of RA projection neurons, that control the activity of vocal motor neurons in the brainstem.

**Fig 4:**
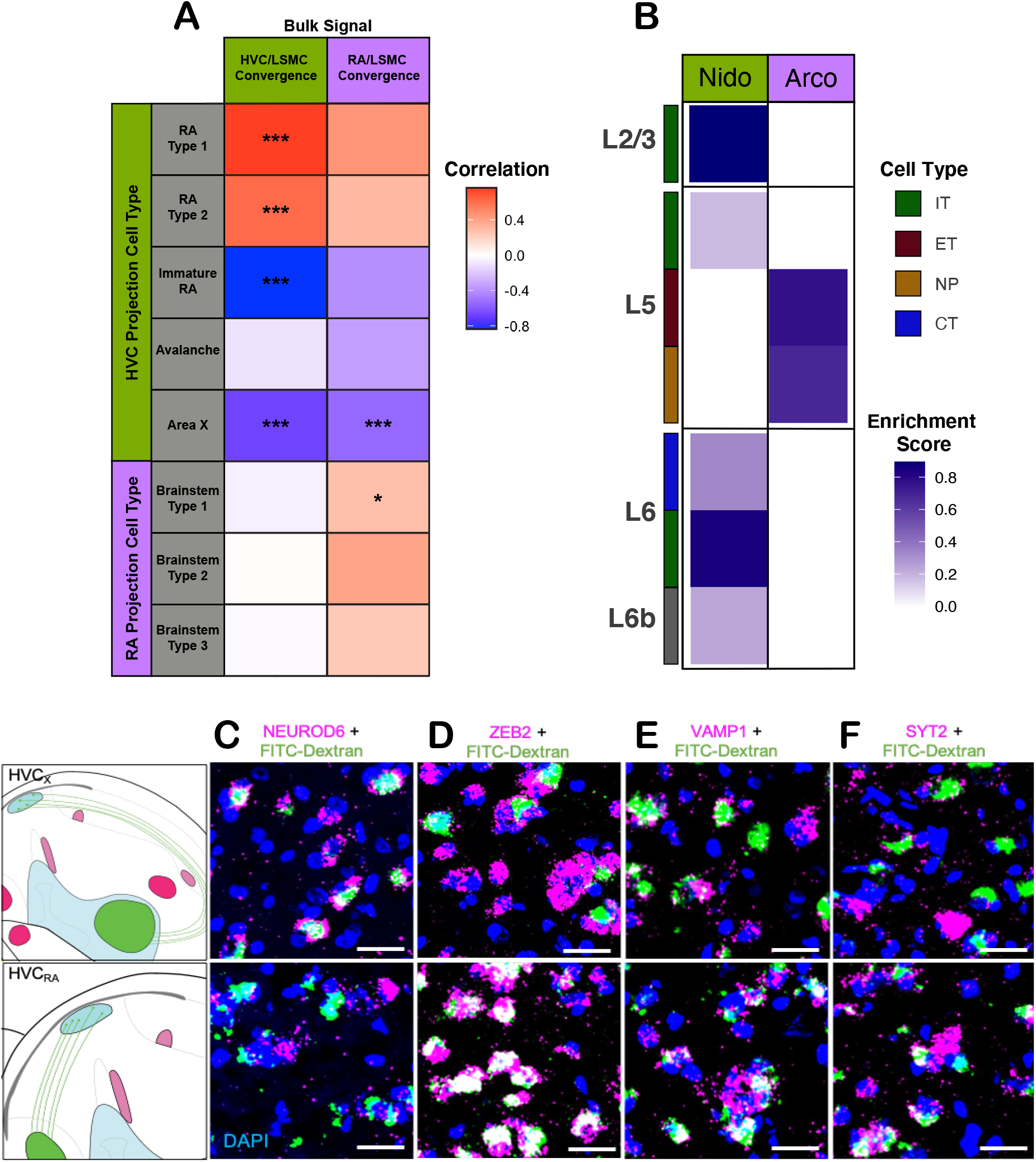
Evidence for convergent microcircuitry for production of learned vocalizations in songbirds and humans. **(A)** Heatmap showing correlations of normalized fold change values between HVC/RA/LSMC convergent molecular specializations (bulk RNA-Seq, this study) and marker genes for glutamatergic cell classes in HVC and RA (snRNA-Seq, Colquitt et al. 2021). Asterisks denote significance at p < 0.05 (*) and p < 0.001 (***). The HVC/LSMC convergent specializations were strongly correlated with markers for HVC_RA_ projection neurons, suggesting these cells are the primary drivers of the observed molecular convergence. **(B)** Heatmap showing enrichment scores of gene markers of human glutamatergic cell types and finch nidopallium (green, HVC parent population) and arcopallium (purple, RA parent population). The intratelencephalically (IT) cells of the human primary motor cortex (PMC) are more nidopallium-like, both in the superficial (2/3) and deep (6) layers. The extratelencephalic cells (ET) are more arcopallium like. **(C-F)** Fluorescent *in situ* hybridization of NEUROD6 **(C)**, ZEB2 **(D)**, VAMP1 **(E**), and SYT2 **(F)** expression in the zebra finch HVC. Retrograde labeling of HVC_RA_ neurons (green) and target gene signal (pink) show colocalization (yellow), suggesting this cell type is the principal driver of the observed molecular convergence. Scale bar is 200μM.

Next, we asked which mammalian cortical cell type most closely resembled the brain subdivisions to which songbird HVC and RA belong, the surrounding nidopallium and arcopallium respectively. We utilized a recently published single-nuclei RNA-Seq dataset from the human primary motor cortex (M1)(*12*), and performed a gene set enrichment analysis between zebra finch nidopallium and arcopallium marker genes with each potential cell type analog (Methods). We found that the zebra finch nidopallium most closely resembled intratelencephalic (IT) cells in layers 2/3 and 6 of the human M1, while the zebra finch arcopallium most closely matched extratelencephalic (ET) cells in layer 5 of M1 (**Fig. 4B**), supporting a “nuclear to cortex-layered hypothesis” of avian mammalian brain homologous (*4, 27*).

Finally, we sought to validate these results using *in situ* hybridization of several previously uncharacterized HVC/RA-LSMC convergent marker genes with either single labeling (n = 22) or double-labeling with retrograde tracing of HVC_RA_ neurons by FITC-Dextran injections into RA (n = 4). We confirmed the specialized expression of these gene markers (**Figs. S12-14**), with all co-labeled genes tested showing specialization to the HVC_RA_ cell class (**Fig. 4D-G**). The collective data suggests evolution of a convergent microcircuit made up of analogous projection neuron types that enable the production of learned vocalizations in songbirds and humans.

### Function of convergent specialized genes in song and speech brain regions

If these analogous cell types are important for establishing a specialized convergent vocal learning microcircuit, we expect their gene expression profiles to be significantly enriched for functional categories that help distinguish these song and speech brain regions from the surrounding tissue, such as neuronal connectivity and cell signaling. To test this hypothesis, we performed gene ontology enrichment analysis on the convergent gene set shared between human LSMC regions and either songbird HVC (n = 251) or RA (n = 155). We found significant enrichments in biological processes related to circuit development, including axon guidance, trans-synaptic signaling, and synapse organization (**Fig. 5A, Table S7**). These specializations were confined to neuronal signaling compartments, with signaling channels and receptors exhibiting specialized expression, including voltage-gated ion channels and glutamate receptors. Interestingly, all of the strongest functional enrichments were observed among downregulated genes (**Fig. 5A**), suggesting downregulation can be a powerful driver of functional diversity from neighboring neuronal circuits. This phenomenon was reversed in the striatum, with all functional enrichments localized to the upregulated convergent gene sets (**Fig. 5B**).

**Fig 5:**
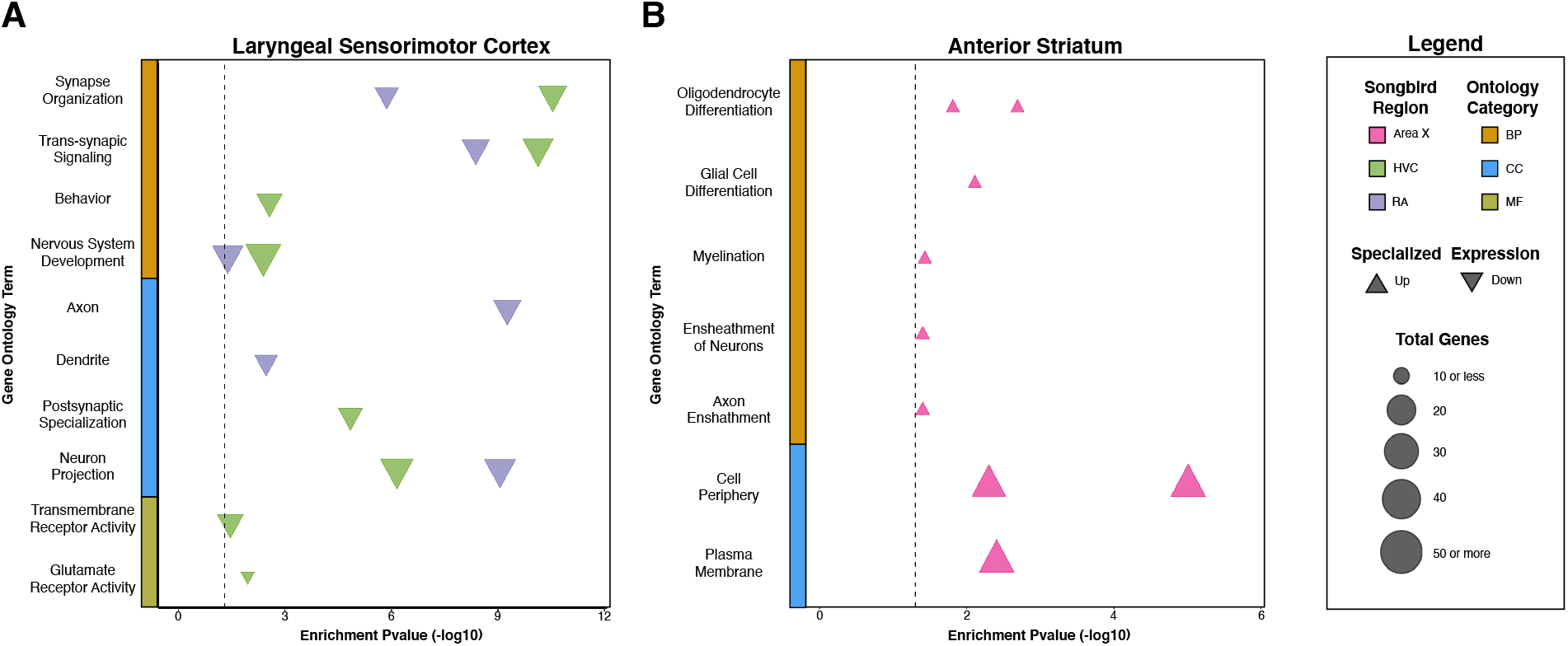
Functional enrichments for genes with convergent specialized expression in songbird and human. (**A**) Gene ontology enrichment plot of convergent HVC/RA and LSMC genes. (**B**) Enrichment plot for convergent Area X and ASt genes. Upregulated and downregulated gene sets from each nucleus were tested separately. Terms shown were significant at FDR < 0.05 (dotted line). Shapes are color coded by specialized songbird nucleus. Direction of shape denotes direction of specialized expression. Size of shape denotes the number of genes contributing to the given GO Term. Genes exhibiting convergent downregulation in the cortex/pallium were functionally enriched, while the opposite phenomenon was observed in genes exhibiting convergent upregulation in the striatum.

One prediction from these findings would be that disruption of these convergent molecular specializations would impact speech function. We cross-referenced our convergent gene sets with genes with known mutations resulting in dysarthria, ataxia, or developmental delay of speech according to the Human Phenotype Ontology database (*29*). We found a significant enrichment for genes involved in speech motor dysfunction in the convergent gene sets of HVC/RA/LSMC (p = 0.02, O/E = 1.46) and Area X/ASt (p = 0.045, O/E = 1.61), suggesting further functional convergence for learned song production (**Table 1**). The identified genes are involved in a range of functions, including intracellular signaling cascades, synaptic transmission, and transcription regulation. All impairments are either the result of single point mutations or large duplication/deletion events, which would disrupt the gene’s protein function or expression. The effects of the human mutations were largely reciprocal of our functional enrichment analyses, with downregulated genes exhibiting gain-of-function phenotypes and upregulated genes exhibiting loss-of-function phenotypes. These data highlight the critical role that many of these convergent genes play in normative human speech and emphasize the value of the songbird as a model to further study their mechanism of action.

**Table 1:**
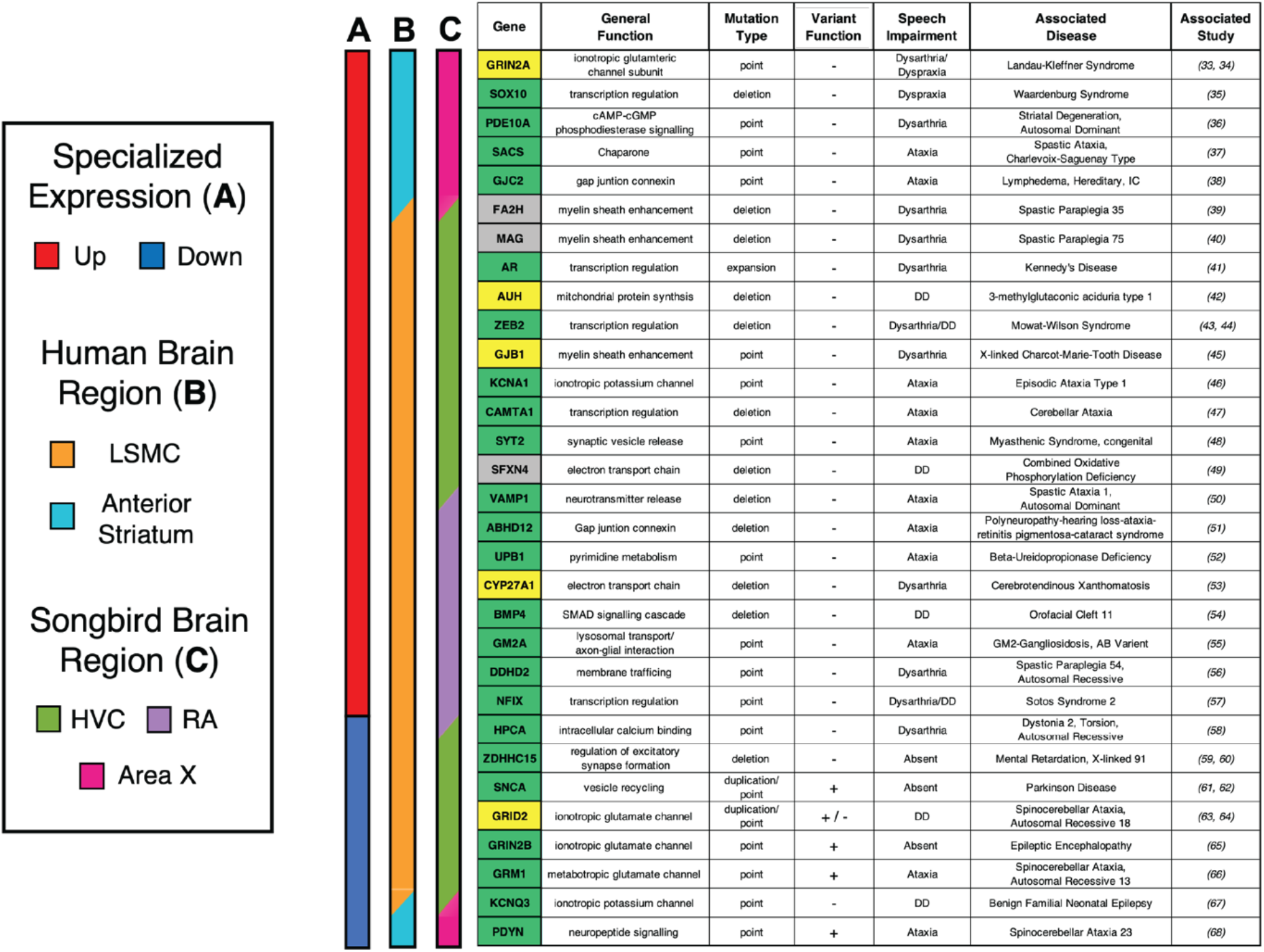
Mutations in genes exhibiting convergent expression specializations in songbird and human song/speech brain regions are associated with human speech dysfunction. List of 32 genes exhibiting convergent expression specializations in songbird and human whose mutation results in delayed/impaired speech phenotypes. Side color bars indicate: (**A**) specialized upregulation (red) or downregulation (blue); (**B**) human specialized expression in LSMC (orange) or anterior striatum (cyan); and (**C**) songbird specialized expression in Area X (pink), HVC (green), or RA (purple). Genes with multiple colors in these bars (slashes) indicate specialized expression in multiple regions. Genes with validated expression via in situ hybridization in the songbird are noted as true positive (bright green), inconclusive (yellow), or not tested (grey).

## Discussion

In the present study, we identify previously unknown patterns of molecular convergence in songbird and human brain regions controlling learned vocalizations of song and speech. We found robust molecular convergence between songbird HVC/RA and human LSMC, with human dLMC exhibiting the strongest match (**Fig. 6**). We did not find a robust molecular convergence between the LMAN song nucleus and human Broca’s area, in contrast to a prior hypothesis that proposed their functional analogies (*7*). The lack of molecular similarity between LMAN and Broca’s area could mean that these brain regions are defined by more unique gene expression profiles in each species. However, further investigation is necessary to confirm molecular and functional divergence, including the in-depth sampling of rare projection neuron classes in Broca’s area with single-nuclei RNA-seq, as well as generating datasets for other more advanced vocal learning species with LMAN analogs, like the parrot NAO and hummingbird VAN nuclei.

**Fig 6:**
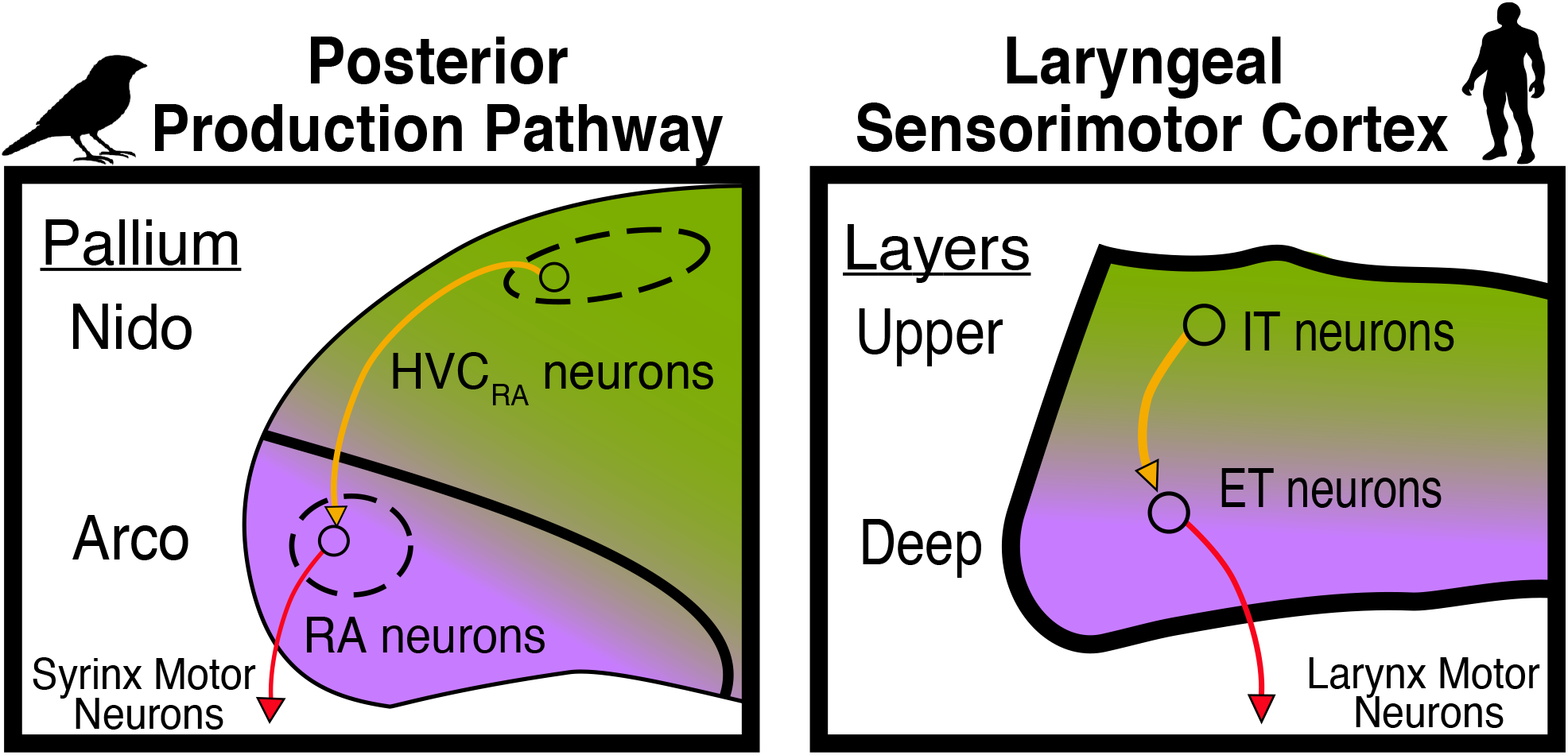
Molecular convergence supports parallel microcircuitry between songbird HVC/RA and human LSMC. Projection neurons in songbird posterior forebrain pathway nuclei HVC (green) and RA (purple) exhibit analogous microcircuitry with human intratelencephalic (IT) neurons (orange arrows) and extratelencephalic (ET) neurons (red arrows) in the human LSMC respectively. This circuit convergence is hallmarked with robust downregulation of neuronal signaling and axon guidance genes. There is additional molecular and circuit convergence between songbird Area X and the human anterior striatum (not shown).

The avian PFP (HVC and RA) is spatially distinct in the songbird brain and controls the production of learned vocalizations through a class of specialized projection neurons. Our results indicate the existence of a molecularly analogous group of projection neurons in the dLMC cortical column, suggesting the evolution of a convergent microcircuit in both species. This is consistent with the functional findings that apparent projection neurons of both songbird RA and human dLMC modulate pitch through direct control of vocal motor neurons in the brainstem, a key characteristics of vocal learners (*20, 30*). Further highlighting their importance for premotor/motor coordination of speech, are their specialized genes associated with speech production deficits when mutated in humans. Whether these genes control similar aspects of vocal production learning in songbirds is an open question. Future work to alter the expression of specialized genes should help provide insight on shared molecular mechanisms underlying song and speech. A shared phenotype following disrupted expression in both species would offer strong evidence that molecular convergence can give rise to similar microcircuitry and functional convergence, even in brains with vastly different global cortical organization.

Indeed, the shared molecular specializations described here further elevate songbirds as an invaluable model system to study the function of vocal motor systems in humans and other vocal learning species. The specialized downregulation of axon guidance gene families like ephrin (*EPH4-6*), plexin (*PLXNA1,C1*), and slit (*SLIT1,3*) suggest that modifying the expression levels of these and other genes could be a powerful mechanism of selectively targeting different regions in the cacophony of cortical gene expression. Convergent positive selection in regulatory elements of surrounding transcription factors exhibiting convergent downregulation in avian and human vocal learners, like *NEUROD6*, implies a shared evolutionary mechanism for developing vocal learning circuitry (Cahill *et al*., *2021*). The *ZEB2* transcription factor, a convergent upregulated gene in songbird HVC and human LSMC, has known roles in axon guidance, and no current animal model captures the striking speech deficits associated with its mutation in patients with Mowat-Wilson Syndrome (Srivatsa *et al*., 2015). Future experiments modulating or even reversing this specialized expression in the songbird vocal motor system will broaden our understanding of the convergent molecular mechanisms for vocal learning circuit dynamics across species.

Molecular convergence governing behavioral convergence for vocal imitation in songbirds and humans highlights important evolutionary constraints on vertebrate motor circuits. We interpret these findings to indicate that vocal imitation maybe an emergent property from the evolutionary modulation of a specific set of genes for neuronal firing and connectivity between brain regions of a vocal motor learning circuit. Songbirds and humans may have evolved vocal imitation due to the modification of these core genes, either through convergent evolution from a deep homology in genes and brain structure (*31*). Regardless of origin, we predict the molecular specializations found here will be present in other vocal learning species, as seen in the analogs of RA for parrots and hummingbirds for the initial set of genes discovered (*4, 32*). However, more extensive screens in these species, as well as other vocal learning mammals (bats, cetaceans, pinnepeds, elephants) will be necessary to determine the extent of molecular convergence for this trait in nature.

Together, our results describe remarkable molecular, circuit, and behavioral convergence to produce learned vocalizations in songbirds and humans. Such parallels open new possibilities for cross-species translation of molecular mechanisms from the songbird to inform the research of human speech dysfunction.

## Acknowledgments

We acknowledge Thomas Carrol and his Bioinformatics team at Rockefeller University for their insightful recommendations to advance this project. We thank Samara Brown for her mentorship and support while conducting this work.

## Data and materials availability

All zebra finch RNA sequencing data generated from this study are deposited in the NCBI SRA archive (BioProject ID: <u>PRJNA678351</u>). All analysis code (zebra finch differential expression, cross species gene set enrichment for microarray and single-cell data) as well as the related data objects (Allen Human Microarray data and zebra finch refseq-ensembl conversion) are available on Google Drive (https://tinyurl.com/GeneConvergence)

## Supplementary Materials

Materials and Methods

Supplementary Text

Figs. S1 to S14

Tables S1 to S7

References (*69-100*)

