## Supplemental Methods and Text for "Convergent gene expression highlights shared vocal motor microcircuitry in songbirds and humans"

**This PDF file includes:**

Materials and Methods
Supplementary Text
Figs. S1 to S14
Captions for Tables S1 to S7

**Other Supplementary Materials for this manuscript include the following:**

Table S1 to S7

### Materials and Methods

Some of the RNA-Seq data generated for this study (surrounding song nuclei) were used and described in a companion study on avian brain organization as described in Gedman et al. (*69*). Several methods are describing in detail in that study, including selection of animal subjects utilized in this study, isolation of brain regions of interest using laser capture microscopy, preparation of samples for RNA-Sequencing, and initial data quality control. In brief, we isolated the four principal song nuclei of the song system (the nidopallial nuclei HVC and LMAN, the arcopallial nucleus RA, and the striatal nucleus Area X) and the surrounding non-vocal motor brain regions for each nucleus as defined according to the activation patterns for non-vocal motor movements (*70*) from a total of four individual adult male zebra finches (**Fig. S1**). Total RNA was isolated from each sample with a minimum RIN of 6 (mean = 7.1) before sequencing library preparation. Following cDNA construction, all samples were sequenced on NextSeq 500 system from Illumina at 150bp paired-end reads with a sequencing depth of ~ 20 million reads per sample. Reads were mapped using Salmon (v0.14.1) to a new high-quality, long-read based, zebra finch reference genome (bTaeGut1_v1, RefSeq Accession: GCF_003957565.1) generated by the Vertebrate Genomes Project (*71*).

#### Zebra finch RefSeq and Ensembl genomic annotations

We downloaded both the RefSeq (Accession: GCF_003957565.1) and Ensembl (bTaeGut1_v1.p, Release 99) versions of the zebra finch transcriptome annotation and loaded both into R with the "makeTxDbFromGFF" function from the GenomicFeatures package (v1.36.4). Both annotation databases were summarized to unique transcripts using the "tidyTranscripts" function from GenomicFeatures package and converted to data frames for easier manipulation. Each annotation had different unique identifiers for chromosomes containing genomic coordinates of gene models. To compare the two annotations, we downloaded the full sequence report from the RefSeq accession, which contains matching chromosome number (Ensembl) with the RefSeq chromosome accession IDs. We used this information to replace all chromosome numbers with RefSeq chromosome IDs in the Ensemble transcript database, allowing for matching gene model coordinates from both annotations to the same assembly. We compared exons that were found in both RefSeq and Ensembl gene models using the "findOverlaps" function from GenomicFeatures with a minimum overlap of 100 base pairs and at least 50% overlap in total sequence length. The RefSeq gene symbols could then be mapped to a unique Ensembl ID, representing the same gene model across annotation versions. If a gene model in one annotation had multiple matches in the other (e.g. Ensembl often called two or more truncated gene models where RefSeq called one full length model), the longest of the matching gene models was taken as the match across versions. This resulted in 16,052 unique RefSeq to Ensembl gene model matches, with a conversion table we placed on Google Drive (https://tinyurl.com/GeneConvergence). Of these shared gene models, we assessed both completeness and accuracy across both annotations. We defined completeness as the total number of exons called to each model in each annotation. For models with the same number of exons in both annotations, we defined accuracy as the total length of each shared exonic feature. Based on these metrics, we decided to utilize the RefSeq annotation for all downstream analyses, while converting to matching Ensembl IDs for cross-species ortholog comparisons.

#### Quality control for unwanted sources of variation and principal component analysis

To assess the effects of unwanted variation on our data, we utilized the scater package (v1.12.2) to identify any non-biological variables that explained a high proportion of the observed variance, such as collection batch and individual bird replicates. Once identified, all spurious variables were addressed with either direct correction using the "removeBatchEffect" function from limma (v3.40.6) or direct inclusion in the linear models for differential expression testing. Effects of removal of unwanted variation were assessed by selecting the 100 most variable genes prior to correction using the "rowVars" R function and assessing clustering of samples after batch correction. Following variance stabilization normalization, we performed principal component analysis (PCA) with the PCAtools package (v1.0.0) using the top 5% of variable genes in the dataset (n = 1081). We also utilized this package to explore the percent variance explained by each PC (scree plot), the correlation of variables of interest to the first four PCs (egiencor plot) and direct comparison of the most important PCs (biplot).

Song nuclei and neural subdivision differential gene expression

We conducted independent differential expression tests for each of the four song nuclei vs. surround comparisons using DESeq2 (v1.24.0). In each case, a linear model was constructed on the raw gene x sample expression matrix with the design "~ collection.batch + region", as these were the largest sources of unwanted and wanted (biological) variation respectively. This allowed for modeling of the observed expression variance between each song nucleus and its adjacent non-vocal motor brain region, while controlling for unwanted collection batch effects. A similar model with the design "~ collection.batch + subdivision" was used to test the difference between the nidopallium and arcopallium neural subdivisions. All gene tests conducted were subjected to independent filtering from DESeq2, as previously described (*69*). Importantly, all genes were subjected to multiple test corrections (Benjamini – Hochberg; $q < 0.05$) to be considered significantly differentially expressed in a given song nucleus comparison. . Results of all differential expression tests were summarized in **Table S1**, with "1" denoting upregulation, "-1" denoting downregulation, and "0" denoting no change relative to the adjacent non-vocal brain region to the given nucleus. Intersections from each column of this table were used as input to the "draw.quintuple.venn" function part of the VennDiagram package (v1.6.20).

Allen Human Brain Atlas microarray data processing

Normalized data and associated metadata were downloaded from the Allen Brian Atlas microarray data portal for all six human individuals (https://human.brain-map.org/static/download). For each individual, each sample was given a unique ID which consisted of brain region, brain hemisphere, individual of origin, and sample replicate number (i.e. SFG_m_L_10021_2; the second replicate [2] of a region of the medial bank of the superior frontal cortex gyrus [SFG_m] from the left hemisphere [L] of individual 10021). Expression from multiple probes from the same gene were averaged to unique Entrez IDs. Data from all individuals were concatenated into a gene (n = 20,788) x sample (n = 3681) data matrix for further processing. To facilitate cross-species comparisons, all gene entrez IDs were mapped to unique Ensembl IDs using bioMaRt (v2.40.5, Ensembl Release 98), resulting in 18,415 genes in the final expression matrix. Gene (row) and sample (column) metadata were added to the final expression object, as well as the sample hierarchies utilized previously (*72*) to enable easier comparison of brain areas. The full Allen data object was utilized for our downstream comparative analyses and can be downloaded from the manuscript GitHub (https://github.com/birdsaplenty45/MolecularConvergence).

Human speech region differential expression analysis

To enable cross-species comparisons, all tests were limited to one-to-one orthologs between songbird and human (n=11,475) as determined by bioMaRt (v2.40.5, Ensembl Release 104). All differential expression analyses were restricted to the left hemisphere due to more thorough sampling and well-characterized lateralization for language functions. All samples were mapped to their nearest Broadman's Area (BA) using the label4MRI package (v1.2) in R which aided in mapping of regions of speech activation. We utilized several sources to localize samples to anatomical regions associated with speech, including Broadman's Area classification and fMRI activation during vocal production tasks (*72, 73*). All cortical or striatal samples that fell within these activation coordinates and part of known Broadman's distinctions were labeled as speech regions for each differential expression comparison. The whole cortex was used as a background for comparison with cortical speech regions. The nucleus accumbens was used as a background for comparison with striatal speech regions. A linear model was constructed on the normalized gene x sample expression matrix with the design "~ 0 + speech". This allowed for modeling of the observed expression variance in speech regions relative to all non-speech regions. From this model, contrasts were conducted using limma (v 3.40.6) and genes were called as differentially expressed following multiple tests corrections (FDR < 0.1).

Cross-species gene set enrichment analysis

Given the large difference in dynamic range between RNA-Seq and microarray data, we could not directly compare magnitude of common molecular specializations between songbird song motor nuclei and human speech motor brain regions. Instead, we utilized a gene set enrichment analysis (hypergeometric test) to test the probability of finding the magnitude of shared molecular specializations by chance between the two species. The density of the hypergeometric distribution for each comparison was calculated using the "phyper" function in R with the parameters $q$, $m$, $n$, and $k$ where:

$q$ = number of genes showing shared molecular specializations in songbird and human (SampleHits)
$m$ = number of genes differentially expressed in the given human comparison (PopulationHits)
$n$ = number of genes *not* differentially expressed in the given human comparison (PopulationFails)
$k$ = number of genes differentially expressed in the given songbird comparison (SampleSize)

Genes were considered to exhibit shared molecular specializations ($q$) if they were differentially expressed in the same direction in both the given songbird and human brain regions. All p values from the hypergeometric tests for each cross-species comparison were corrected for multiple tests using the "p_adjust" function in R (method = "fdr") and represent the probability of seeing this level of overlap or greater. Fold enrichment was calculated by dividing the observed overlap ($q$) by the overlap expected by chance (($m$/Population)*$k$). Heatmaps of gene sets exhibiting convergent specialized expression between both species were generated using the "heatmap.2" function from the gplots package (v3.0.3). Gene ontology analysis was conducted on all upregulated and downregulated genes for each comparison using the gprofileR2 package (v0.1.9), utilizing all one-to-one orthologous genes as a custom background.

Assignment of human speech disorder phenotypes from convergent gene set

We queried the Human Phenotype Database (*74*) for genes associated with speech motor dysfunction. We limited our search to robust deficits labeled as "dysarthria", "delayed speech and language development", and "neurological speech impairment". One-to-one orthologs from each of the songbird/human convergent gene sets were intersected with this list to obtain speech disorder genes exhibiting cross-species specialized expression to vocal learning brain regions. Manual literature review was conducted for each gene to further validate and describe its impairment in vocal imitation.

I*n Situ* Hybridizations

A list of song system marker genes (n = 142) determined by in-situ hybridization was provided by Lovell and colleagues as part of the Zebra Finch Expression Atlas (*75*) and were used to assess the percent accuracy of our differential expression results. We also generated our own in-situ hybridizations on selected genes (n = 32). All single-label, colorimetric in situs were performed using a method described in Biegler et al. (*76*). To determine overlap of multiple genes in the same cells, we performed double-label fluorescent *in situ* hybridization. Plasmids containing RNA polymerase promoters and cDNA sequences for genes of interest were used to amplify the cDNA inserts by PCR. We also used an alternative method, where we isolated the cDNA insert from the plasmid with *PvuII-HF* (New England Biolabs, Cat. #R3151) or *BssHII* (New England Biolabs, Cat. #R0199) restriction enzymes sites flanking the probe sequence. The cDNA products were purified using the Nucleospin® Gel and PCR Cleanup kit (Takara Bio, Cat. #740609). The cDNAs were transcribed and labeled following the instructions provided with the DIG RNA Labeling mix or the Fluorescein RNA Labeling mix (Sigma, Cat. #11685619910). The generated RNA probes were purified by ethanol precipitation, resuspended in 90% formamide, and stored at -80$^{o}$C until further use.

All steps were performed at room temperature (RT) unless specifically noted. Slides containing 4-6 brain sections of a series were fixed with 4% Paraformaldehyde (PFA) in 1X PBS, washed with 1X PBS and then incubated in acetylation buffer (250mL 0.1M Triethanolamine, 280µL NaOH and 625µL acetic anhydride, mixed right before use). The sections were washed with 1X PBS and dehydrated serially with 70%, 95% and 100% EtOH. Samples were incubated for at least 1 hour in prehybridization solution (50% formamide, 5X SSC, 1X Denhardt's solution, 250µg/mL tRNA, 500µg/mL herring sperm DNA).

For double labeling, we diluted the two probes of interest (one conjugated with DIG and one with FITC) into the same tube of hybridization solution at a ratio of 1:100 for each probe. We then followed the same steps as the single label protocol, stopping after the final 0.1X SSPE wash for 5 min at room temperature. At that point, slides were washed with 1X PBS for 5 minutes, incubated in 10% $H_2O_2$-1X PBS for 10 minutes to remove any endogenous peroxidase activity and then washed twice in 1X PBS for 5 minutes. To label the hybridized DIG-conjugated probes, the sections were incubated in 0.5% Roche Blocking Solution for 1 hour and then incubated with Anti-DIG-POD antibody (Sigma, Cat. #11207733910) at 1:1000 dilution in 0.5% Roche Blocking Solution (Sigma, Cat. #11096176001) overnight at 4°C.

The slides were washed twice with 1X PBS for 10 minutes and then once with 0.1% BSA-1X PBS for 10 minutes. The remaining steps all occurred while protecting the slides from natural or ultraviolet light, utilizing opaque horizontal slide boxes or vertical slide mailers. Slides were incubated with 1:100 Cy3-TSA amplification reagent in 1x Plus Amplification Diluent (Akoya Biosciences, Cat. #NEL741001KT) for 15 minutes, then washed in 1X PBS for 5 minutes. The slides were then quenched in 15% $H_2O_2$-1X PBS for 30 minutes to remove any remaining peroxidase reactivity and washed twice in 1X PBS for 5 minutes. Slides were fixed in 4% PFA-1X PBS for 5 minutes and washed twice with 1X PBS for 3 minutes to ensure tissue integrity.

For the second round of antibody staining against the hybridized FITC-conjugated riboprobe, the sections were incubated in 0.5% Roche Blocking Solution for 30 minutes and then incubated with Anti-FITC-POD antibody (Sigma, Cat. #11426346910) at 1:1000 in 0.5% Roche Blocking Solution for 2 hours at room temperature, or overnight at 4°C. The slides were washed twice with 1X PBS for 10 minutes and then with 0.1% BSA-PBS for 10 minutes. The slides were incubated with 1:100 FITC-TSA in amplification buffer for 15 minutes, then washed in 1X PBS for 5 minutes. The sections were counterstained with DAPI-1X PBS for 15 minutes. The slides were washed twice in 1X PBS for 5 minutes and then rinsed with dH2O. They were then cover-slipped with Prolong Diamond Antifade mounting media (Invitrogen) and dried overnight at room temperature, protected from light.

Comparison with human single nucleus RNA-seq data

We utilized a single nucleus RNA-seq dataset of human primary motor cortex from Bakken and colleagues (*77*). The expression data object and cell type assignments were provided by the authors. The single nuclei expression data was processed and $\log_2$ normalized with the Seurat package (v3.1.1), resulting in expression for 32,333 transcripts from 67,127 single nuclei. Each nucleus was assigned to a broad cell class of either glutamatergic (n = 48,536), GABAergic (n = 23,992), or non-neuronal (n = 4005). Each broad class was further divided into distinct subclasses of glutamatergic (n = 8), GABAergic (n = 6), and non-neuronal (n = 6) cell types. To test for expression similarity to broad zebra finch pallial subdivisions, we first z-score normalized the entire expression matrix. Next, we subset the data for genes that were specialized in the zebra finch nidopallium and arcopallium (relative to each other) and shared one-to-one orthology with human. Using the specializations for each avian neural subdivision, we calculated an enrichment score for each cell in the human dataset with the following formula:

$$\mathbf{E = \Sigma\ (U_{Z\text{-}score}) - \Sigma\ (D_{Z\text{-}score})}$$

Where E is the enrichment score calculated by subtracted the sum of the human Z-scores from all downregulated zebra finch genes (D) from the sum of the human Z-scores from all upregulated zebra finch genes (U). The more positive this enrichment score the more a given cell in the human primary motor cortex exhibited expression bias towards the reference zebra finch subdivision (e.g. nidopallium). We calculated nidopallium and arcopallium enrichment scores for each cell independently. We then selected the glutamatergic cell type subclasses and plotted their median enrichment scores in a heatmap.

Comparison with zebra finch single nucleus RNA-Seq data

In order to assess the principal cell types driving the convergence signal with human LSMC in our bulk expression data, we utilized results from a published analysis of single nuclei RNA-Seq data from zebra finch HVC and RA (*78*). The authors reported cell type specific markers for five principal glutamatergic neurons in HVC and three principal projection neurons in RA, along with $\log_2$ fold changes for each marker. We correlated these fold changes with those of genes that exhibited convergence with LSMC and either HVC or RA from our bulk expression data. If a gene did not appear in both the single nuclei and bulk expression datasets it was removed from the test. We conducted a two-sided test for significant Pearson correlation using the "cor.test" function in

R. All correlations were reported and visualized in a heatmap generated with the “heatmap.2” function from the gplots package (v3.0.3).

### Supplementary Text

Regions of interest, sequencing statistics, and annotation assessments

We were able to map nearly 98% of reads to the new VGP zebra finch assembly, compared to 87% to the previous Sanger-based assembly (*71, 79, 80*), maximizing the power of our analysis. Two gene annotation models were available for the VGP assembly from NCBI (RefSeq Accession: GCF_003957565.1) and Ensembl (Release 98), so we assessed the quality of each for gene model consistency and sample expression quantification. We found that for gene models called over the same genomic coordinates, the RefSeq annotation models contained significantly more exons on average than the Ensembl annotation (**Fig. S2A**). For gene models with the same number of exons, we found that, on average, these exons were significantly longer in the RefSeq annotation than in the Ensembl annotation (**Fig. S2B**). To maximize the number of reads utilized from our experiment, we opted to use the RefSeq annotation for quantification of aligned reads. All sequence reads that uniquely mapped transcripts as defined by the RefSeq annotation were summarized into a gene x sample matrix utilized for all downstream analyses.

However, for later comparison with human gene expression data, we needed to confidently define one-to-one gene orthology between species, a task more easily accomplished with the Ensembl annotation and BioMart query system. To leverage both the superior gene model predictions from RefSeq and the orthology information from Ensembl, we aligned both annotations together to find gene models shared and generated an ID conversion table between versions. We retained all gene models in the same coordinate space with more than 75% overlap in their exon regions, which resulted in 16,010 genes present in both RefSeq and Ensembl (72% of all possible genes, 91% of possible protein coding genes). We utilized this list to convert RefSeq gene symbols to unique Ensembl IDs for comparison to human in later analyses.

Data quality control and sources of unwanted variation

To maximize the effects from biological variance, we first tested whether there were any batch affects or other covariate influences on the gene expression patterns across all samples. We generated a density plot correlating gene expression variance to variables of interest, with peak size indicative of number of genes effected (**Fig. S3A**). We found that sample group (blue), which included broad brain tissues (pallium vs. striatum) and their subregion distinctions (nucleus or surround) was a broad peak with many genes that explained on average 42% of the variance across all samples. The second largest variable effecting a comparable number of genes was sample collection batch (yellow), which explained on average 25% of the variance. The effect from individual replicate (green) explained a comparable proportion of the variance (20%) but effected fewer genes, which is expected with high interindividual variability and consistent with previous analyses of a subset of this dataset (*69*). Other covariates, like library batch (red), RNA concentration (purple), and RNA quality (brown) had a relatively small impact on the data (6-13% of the variance explained). The potential explanation of this large collection batch effect is that, in addition to its own variance, this variable is confounded with the individual effect, as we did not randomize sample collection and RNA isolation across the batches. We were able to model and remove the effects of collection and library batch and using limma "removeBatcheffect" to generate a covariate-corrected count matrix. This correction reduced the number of genes effected by collection batch (**Fig. S3B**). We visualized the effect of removing sources of unwanted variation by plotting the expression of the top 100 most variable genes before

(**Fig. S3C**) and after (**Fig. S3D**) covariate correction. Prior to correction, some samples clustered more by individual replicate (bird 1) than by biological replicate (**Fig. S3C,** Area X & RA**)** and this effect was removed after covariate correction (**Fig. S3D**).

To assess the principal sources of biological variation in our covariate-corrected data, we performed PCA using the top 5% of most variable genes (n=1081). We found that the first four PCs explained 78% of the variance in the experiment (**Fig. S4A**). We found strong correlations for each of these principal components to variables of interest (**Fig. S4B**). PC1 (43.93% variance) was highly correlated ($R^2 = 0.98$; $p < 0.001$) with brain subdivision and explained the difference between striatum and the rest of the pallium samples. The negative correlation to RNA concentration ($R^2 = -0.61$; $p < 0.001$) can be explained by the larger sample area taken from the striatum and associated Area X (**Fig. S1**). PC2 (17.23% variance) and PC3 (10.51% variance) were significantly correlated ($R^2 = -0.47, 0.64$; $p < 0.01, 0.001$ respectively) with the vocal nuclei and best explained the difference between song system and the surrounding samples. PC4 (6.67% variance) showed a significant anticorrelation ($R^2 = -0.37$; $p < 0.05$) with the vocal nuclei. Consistent with these correlations, a PCA plot of PC1 vs. PC3 best shows the separation of the samples by cell population and song nuclei specializations (**Fig. S4C**). These results demonstrate robust signals of biological variation in our expression data that improve when controlling for sources of unwanted variation. We included collection/library batch as covariates in the linear models for differential expression where possible.

<u>Validation of bioinformatic results with in situ hybridizations</u>

To assess the accuracy of the genes called differentially expressed in the zebra finch song system (**Fig. 2**), we conducted in situ hybridization analyses of 178 genes. These genes were selected based on high values in several metrics, including number of specialized nuclei, magnitude of fold difference, strength of q-value, and availability of data from other sources. Genes were called either true positive (i.e. agrees with DE call), false positive (i.e. disagrees with DE call), true negative (i.e. agrees with non-DE call), or false negative (i.e. disagrees with non-DE call). These classifications were determined by independent blind assessment from authors MB and MW. We conducted this assessment for general differentially expressed genes in the zebra finch (**Fig. S5**), genes found to exhibit convergent differential expression with human (**Fig. S12**), as well as convergent genes associated with speech disorders from the human LSMC (**Fig. S13**) and anterior striatum (**Fig. S14**).

<u>Convergent specializations between songbird and human vocal learning brain regions</u>

To test for molecular convergence between zebra finch song nuclei and any of the hypothesized analogous human speech motor regions, we utilized the same publicly available human brain microarray dataset from the Allen Brain Institute (*81*) used in past studies of vocal learning molecular convergence (*72, 81*). After data cleaning, the final human dataset consisted of 18,415 genes with unique Ensembl IDs, across 3,681 brain regions from 6 individuals (**Table S3**). Each sample is associated with specific MNI-coordinates, allowing for mapping of speech motor regions based on anatomical (Broadman's Areas) or functional (fMRI activation) designations from the literature. Utilizing this coordinate space, we investigated the molecular specializations in brain regions shown to be active during speech production and learning (*73, 82, 83*). These included five cortical (dLMC, dLSC, vLMC [collectively LSMC], SMA, and Broca's area), and two striatal (anterior caudate and putamen) regions with samples restricted to activation coordinates (**Fig. S6**). We found the genes marking these brain regions to be largely distinct, apart from the LSMC and anterior striatum, which shared a high degree of overlap in their specialized expression (**Fig. S7**).

For the gene set enrichment analysis using a hypergeometric test for significant overlap in cross species specialized expression (**Fig. S8**), we first defined the universe for all comparisons as all one-to-one orthologs in both species (n=11,475) as defined by Ensembl (Release 104). We tested molecular specializations of each zebra finch song nucleus (set 1) with each human speech motor area (set 2) and determined the size of their shared gene set. Importantly, a gene had to exhibit specialized expression in the same direction (e.g., upregulated) in both species to be counted in the shared set. For each shared gene set, we calculated an enrichment p-value denoting the probability of finding the observed overlap or greater from the starting species-specific gene sets. Any significant overlap (FDR < 0.05) in specialized expression was interpreted to represent convergent specialization in the given vocal learning brain regions. The results of each comparison are described in detail in the subsequent sections.

Laryngeal sensorimotor cortex (LSMC)
See Main Text.

Anterior Striatum
The striatum is a region of the basal ganglia with a well-established role in the organization and generation of voluntary movements across vertebrates (*84*). The human striatum is composed of three primary subdivisions: the caudate, putamen, and nucleus accumbens. The caudate and putamen are the primary sites of cortical inputs conveying information on the desired movements to execute. Individuals with Parkinson's Disease exhibit degenerated midbrain dopaminergic neurons that project to the striatum and as a result experience the involuntary inability to initiate movements, including speech.

Several studies have highlighted the functional and connectivity specializations of the striatum for speech production. Functional MRI studies have shown the anterior portion of the striatum is active for volitional production of learned speech sounds (*82, 83*). Functional connectivity studies have shown the anterior caudate and putamen receive inputs from the SMA and the Area6v in monkeys and dLMC in humans, forming a critical junction in a speech motor cortico-striatal-thalamic loop (*85, 86*). The putamen receives more robust innervation from these areas and therefore might be more important for learned speech production (*87, 88*). Consistent with this idea, lesions to human putamen result in impairments in many aspects of learned vocal motor control (e.g. production, imitation, intonation etc.), while lesions to non-human primate putamen have little effect on innate monkey vocalizations (*89*).

The songbird striatal nucleus Area X exhibits similar connectivity, functional, and molecular parallels to the human anterior caudate and putamen (*83, 90*). Both Area X and human striatum are critical components of a forebrain-striatal loop necessary for learning and execution of motor skills. Lesions to Area X in adult birds results in impair song production with a stuttering-like phenotype, suggesting a role in organization of learned vocalization sequences (*91*). Mutations in the human FOXP2 transcription factor, prominently expressed in the striatum, result in impaired speech acquisition and production (*92*). This gene is also down-regulated in songbird Area X in response to singing behavior (*93*), and FOXP2 knockdown animals show incomplete and inaccurate imitation of tutor song (*94*). These molecular parallels extend beyond a single molecule, as one of our previous studies showed that Area X exhibits convergent molecular specializations with the human anterior caudate and putamen, with a stronger association to the later (*72*). However, due to limitations of the songbird microarrays, there might be additional genes part of this convergence signal that could inform shared molecular mechanisms of function in both species.

We tested for convergence in molecular specialization between songbird Area X and portions of the human anterior caudate/putamen active during speech (**Fig. 3D**). We found a significant overlap in specialized expression between Area X and anterior putamen, supporting the findings from Pfenning et al. (*72*). However, we found many more genes (n=117) exhibiting convergent expression between Area X and the anterior putamen than was previously reported (n=78), highlighting the advantages provided of whole transcriptome sequencing. In addition, we found molecular convergence between songbird Area X and the head of the human anterior caudate (n=86), an association that previously did not reach significance. This was driven by many of the same genes in both regions, as the caudate shared nearly all its specialized genes with the putamen (n=82, 95%), suggesting both regions utilize similar genes for speech production (**Fig. 3B**). Importantly, we did not find a significant overlap in specialized expression between these human striatal regions and any other songbird nucleus, or between Area X and any other human non-vocal motor striatal region.

Supplementary Motor Area (SMA)

The human SMA is an important premotor region involved in the initiation and temporal coordination of speech (*95*). This region is composed of two subregions with a functional gradient, with the more posterior SMA proper activated prior to the onset of speech motor movements, and the more anterior pre-SMA active for more complex language tasks (i.e. lexical disambiguation) outside of the motor domain (*96*). Despite this regional specification, the region is often collectively referred to as SMA, and damage to both subregions results in speech disfluency (*97*).

We found a significant enrichment between the molecular specializations of songbird HVC and LMAN and those of the human SMA (**Fig. 3C**). Their overlap was weaker (n=7 and 12 genes, respectively) compared to the LSMC (**Fig. 3A**), but still significant when considering the size of the SMA specialization compared to the rest of the cortex (n=27, FDR < 0.1). Interestingly, we observed a significant overlap of similar strength between these premotor nuclei and the premotor cortex generally, suggesting they are some broad expression similarities with no specificity to the vocal motor domain. Therefore, we did not consider this a meaningful signal of molecular convergence for vocal learning.

Broca's area

Historically, Broca's region has been considered to consist of two subregions in the inferior frontal gyrus (IFG) known as Brodmann's area (BA) 44 (triangular part) and BA 45 (opercular part). Due to the size of the original lesion described in Broca's aphasic patients (*98*), neighboring areas like the more anterior portion of the IFG area 47 (orbital part) may also be considered part of this region. Indeed, a large body of work demonstrates that each of these subregions have selective activation for a number of complex tasks, including (but not limited to) acquisition of grammar rules, speech production, estimation of time intervals, and movement imitation (*99*). A recent study using in-vivo electrophysiology on human patients and found that Broca's area is most active and associated with planning speech production seconds before speech is produced, and more so for speech production social interactions (*100*).

Like the SMA result, we found a significant convergence in molecular specializations between the songbird HVC and Broca's area, as well as the premotor cortex (BA6) generally (**Fig. 3C**). Here, we defined Broca's region as samples from the Allen Brain Atlas whose MNI coordinates fell within the classical BA 44 and 45 boundaries of the IFG. Broca's area (BA 44 and 45) exhibited 50 genes specialized relative to the whole cortex, with HVC exhibiting convergent expression with a small number of these genes (**Fig. S11**, n = 10, 20%). With such small, non-specific overlap in gene expression, we did not consider this a robust signal of vocal learning molecular convergence.

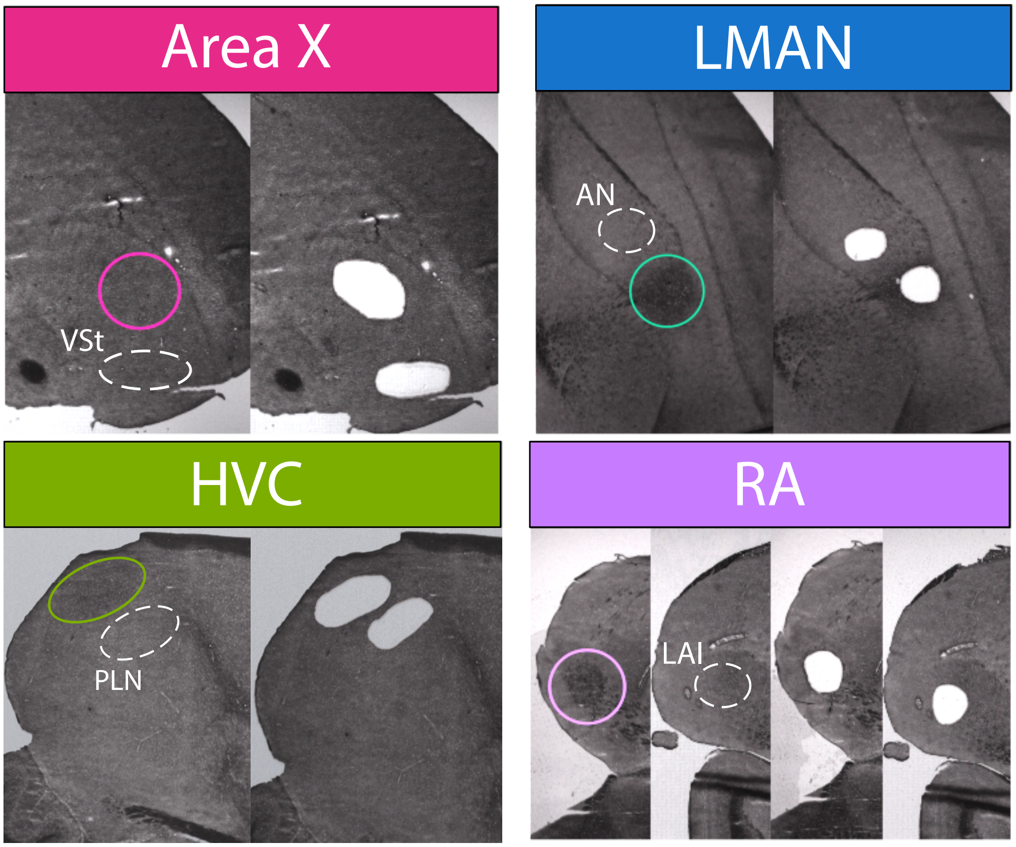


**Figure S1: Example song nuclei and surrounding region isolations with LCM.**

(**A-D**) Brightfield sections showing the brain regions profiled before (left column) and after (center column) LCM dissections. Darker brain regions are due to increased myelination, some of which separate brain subdivisions via axon tracts. Specific song nuclei are highlighted with colored circles and corresponding surrounding pallium are highlighted with dashed white circles. All sections are sagittal, where dorsal is up, and anterior to the right. The LAI section is more lateral to the RA section. VSt: Ventral Striatum. AN: Anterior Nidopallium. PLN: Posterior Lateral Nidopallium. LAI: Lateral Intermediate Arcopallium.

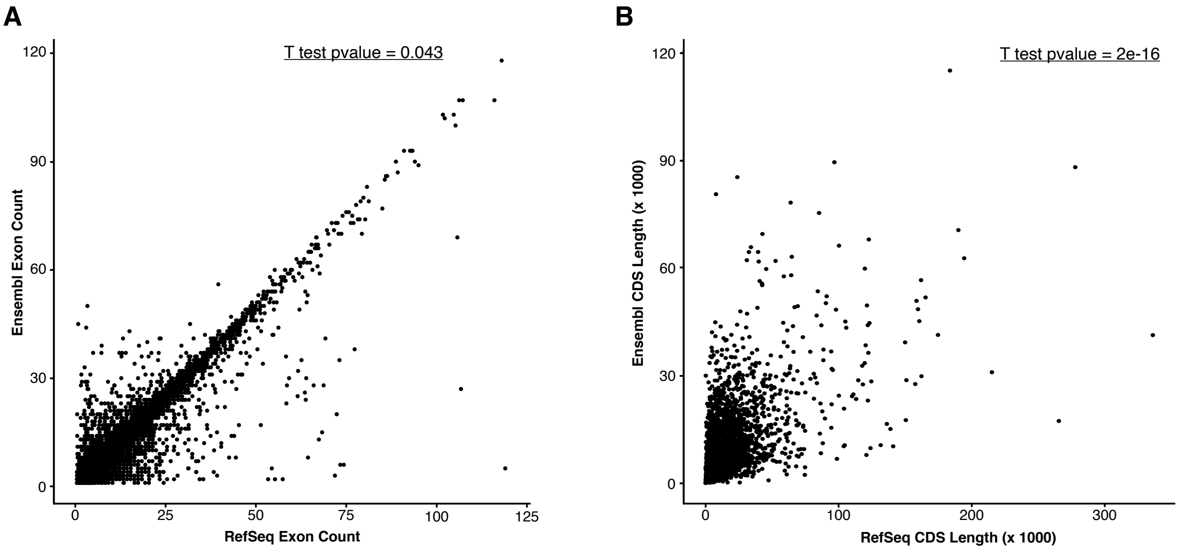


**Figure S2: RefSeq gene models outperforms Ensembl gene models in total exon number and length. (A)** Number of exons defined in gene models sharing the same coordinates (>75% overlap) in RefSeq and Ensembl, with a significant shift towards RefSeq (t-test p-value = 0.043). **(B)** Total gene length for models with the same number of exons called between Refseq and Ensembl annotations. RefSeq genes are significantly longer compared to Ensembl (t-test p-value = 2e-16).

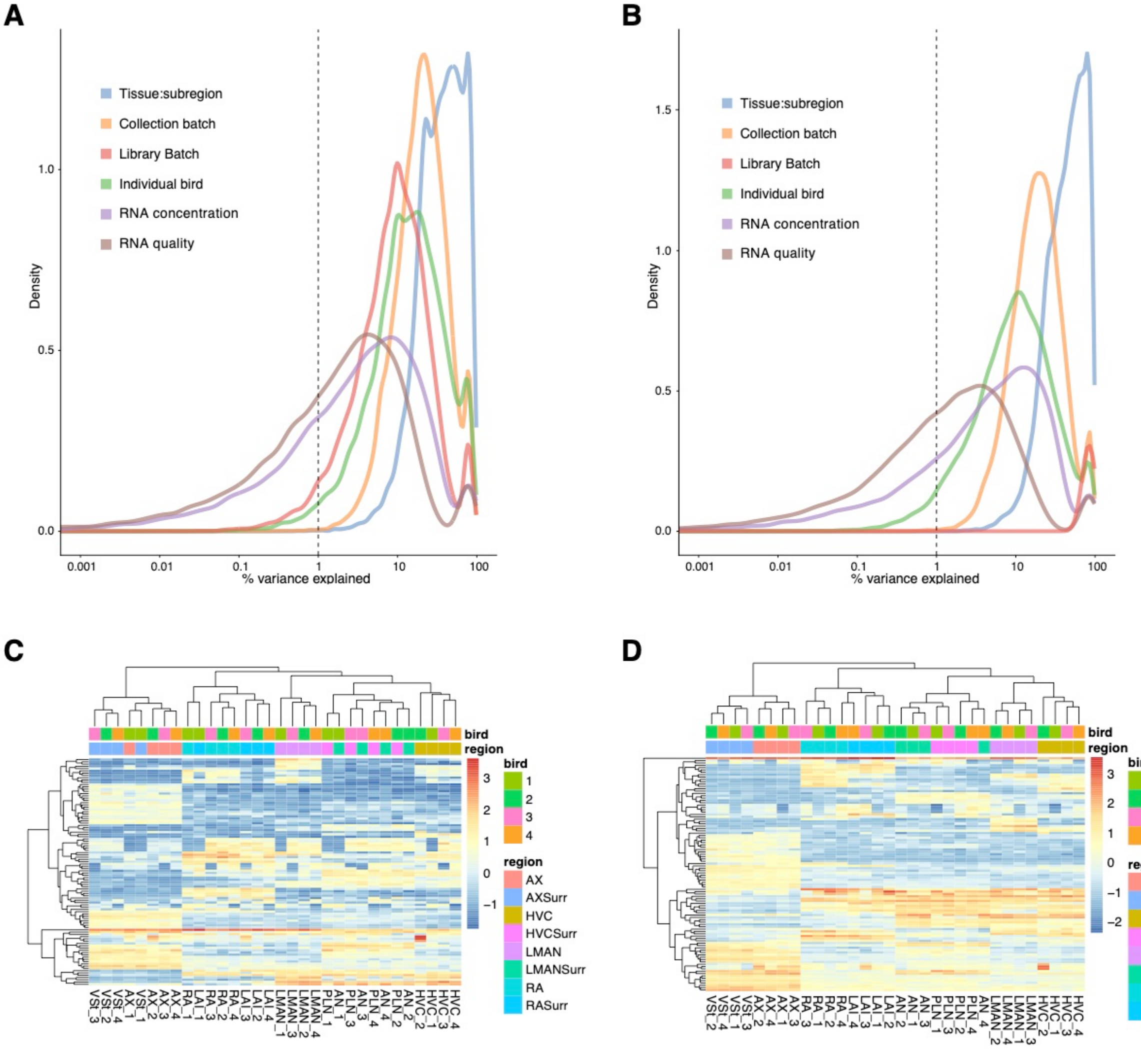


**Figure S3: Exploring and removing sources of unwanted variation.** (**A-B**) Density plots attributing the percentage of variance explained to each variable measured in the experiment. The area under the curve is proportional to number of genes effected. (**A**) Distribution of variance explained by each variable in the normalized data before covariate correction. **(B)** Distribution of variance explained by each variable in the normalized data after covariate correction. The effect of library batch disappears, and the impact of collection batch is greatly reduced. **(C-D)** Heatmap of the top 100 most variable genes in the dataset before **(C)** and after **(D)** correction for unwanted covariates. Prior to correction, a subset of samples clustered more by individual (e.g. Area X and RA of bird 1). Since these samples were collected in the same batch, adjusting for this covariate (**D**) resulted in more robust clustering by anatomical region.

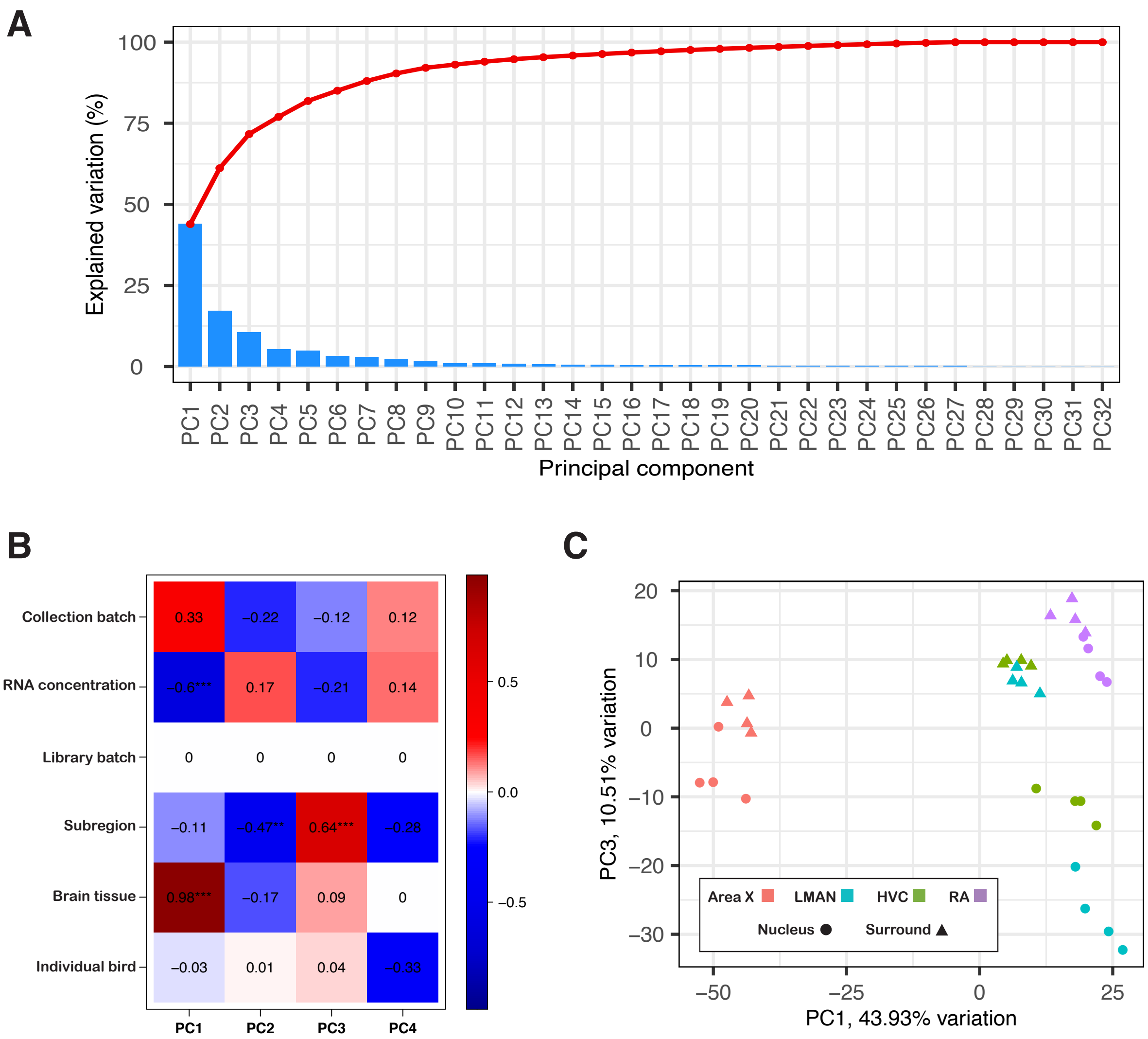


**Figure S4: Principal component analysis of normalized expression data. (A)** Screeplot of percentage of variance explained by each principal component (PC). The number of PCs matches the number of samples in the experiment (n=32). The first four PCs explain nearly 80% of the observed variance. **(B)** Correlation heatmap of variables of interest to the first four PCs. The first PC is best explained by broad brain tissue (pallium vs. striatum), while the third PC is best explained by subregion (song nucleus vs. surround). **(C)** Direct comparison of the most biologically relevant PCs as determined in (**B**). These two PCs (PC1 and PC3) explain 54% of the total variance, with a clear separation of the song nuclei from their surrounding brain regions.

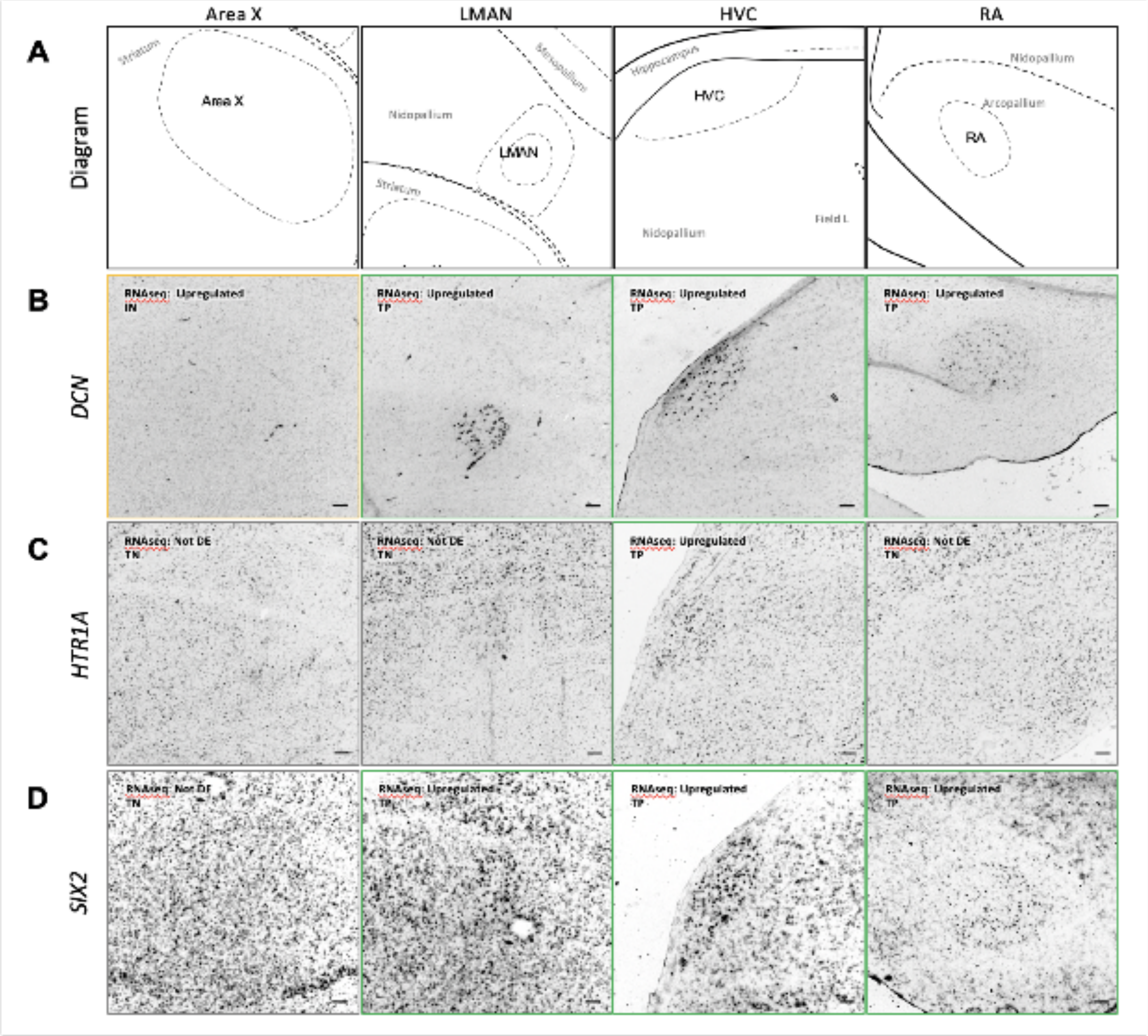


**Figure S5: Validation of zebra finch song nuclei DEGs**. (**A**) Approximate anatomical diagram of adult male zebra finch vocal nuclei. (**B-D**) Colorimetric *in situ* hybridization characterizing (**B**) *DCN*, (**C**) *HTR1A*, and (**D**) *SIX2*. Image outlines are color-coded according to comparison with RNA-seq data. True Positive (TP) = Green; True Negative (TN) = Grey; False Positive (FP) = Red; False Negative (FN) = Blue; Inconclusive (IN) = Yellow. Scale bar = 100μm.

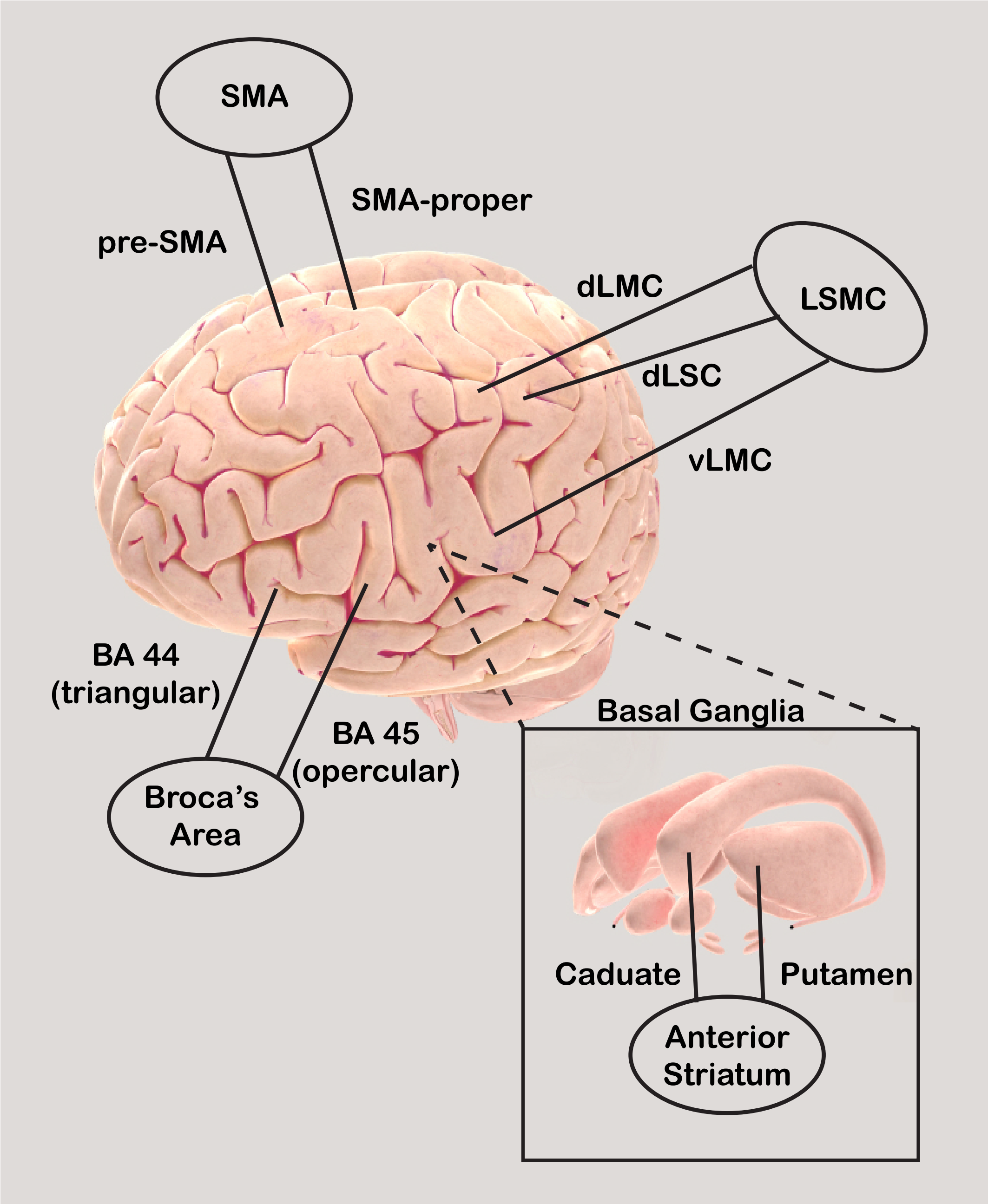


**Figure S6: Anatomical map of human speech related brain regions selected for differential expression testing.** The broad functional area is enclosed in circles and their subregions are pointed to by lines. LSMC: Laryngeal sensorimotor cortex, dLMC: dorsal laryngeal motor cortex, dLSC: dorsal laryngeal sensory cortex, vLMC: ventral laryngeal motor cortex. SMA: supplementary motor area. BA: Brodmann's area.

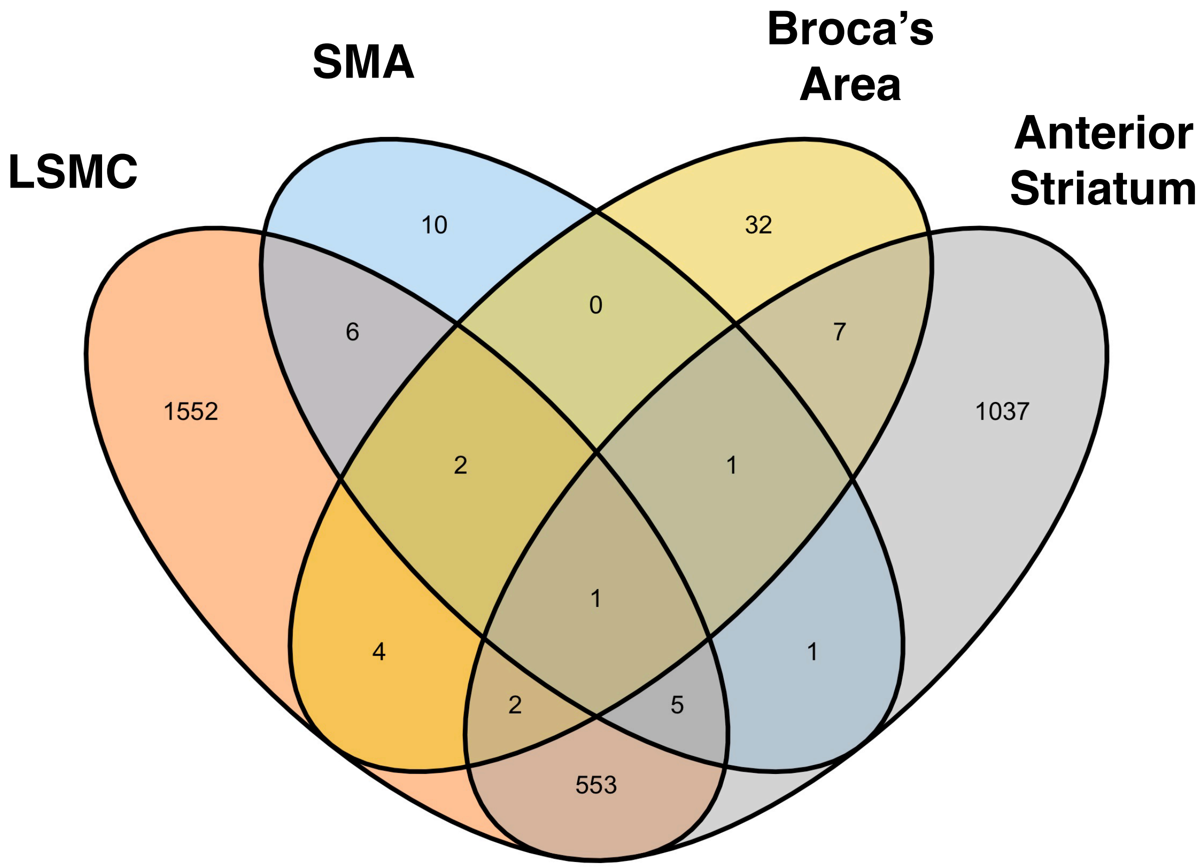


**Figure S7: Shared expression specializations of key nodes in the human vocal motor pathway.** Venn diagram quantifying the overlap in specialized gene expression among the cortical (LSMC, SMA, Broca's) and subcortical (anterior caudate/putamen) speech brain regions. Each region is defined by largely distinct molecular specializations, apart from the LSMC and anterior striatum.

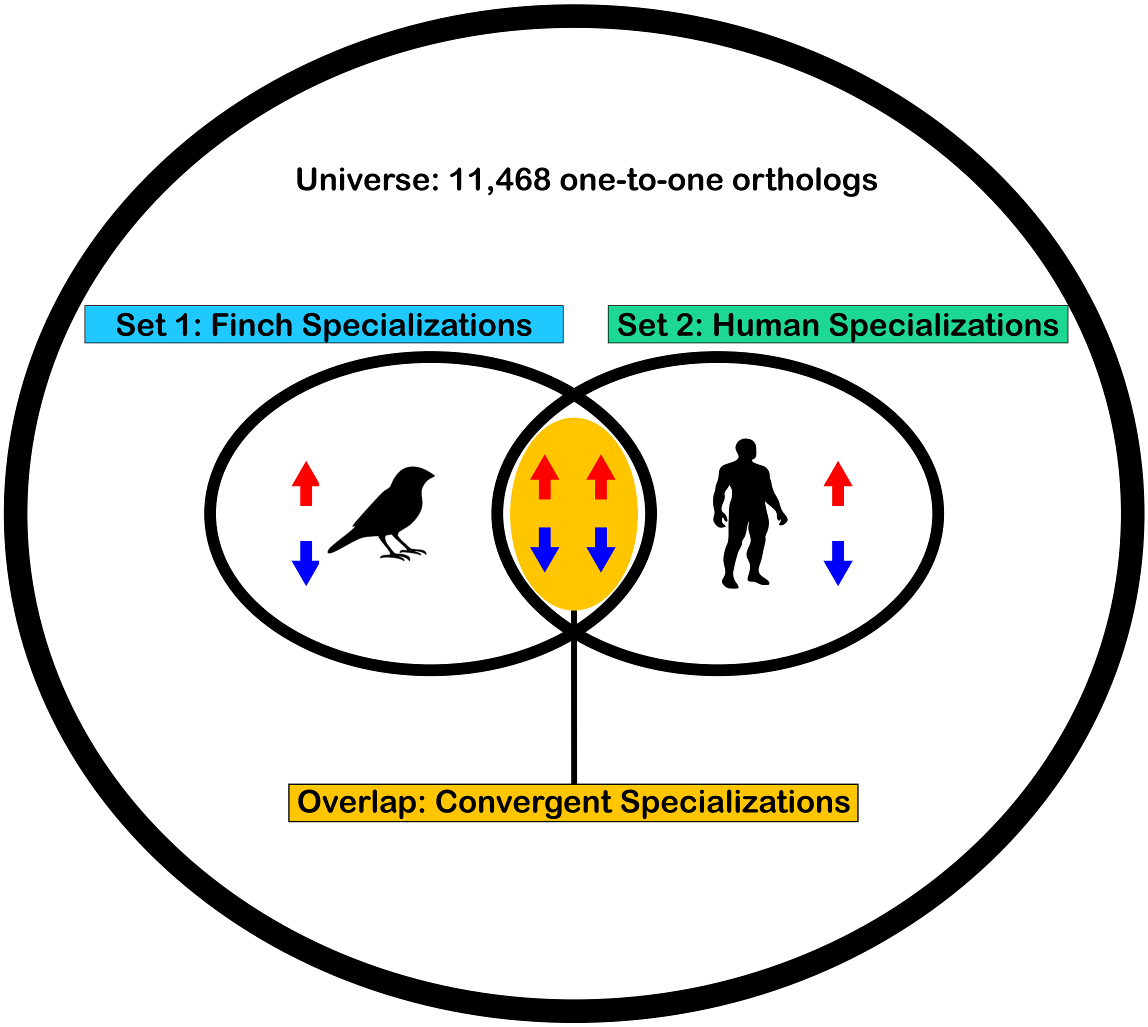


**Figure S8: Hypergeometric test for convergent molecular specializations.** Gene set comparisons were limited to one-to-one orthologs in zebra finch and human (n=11,468). Gene sets from zebra finch (set 1) and human (set 2) were compared for shared specializations with similar regulation. Overlap in specialized genes was quantified and tested for significant enrichment in both species. Shared gene sets determined significant (FDR < 0.5) were considered candidates of convergent molecular specializations for song/speech brain regions in both species.

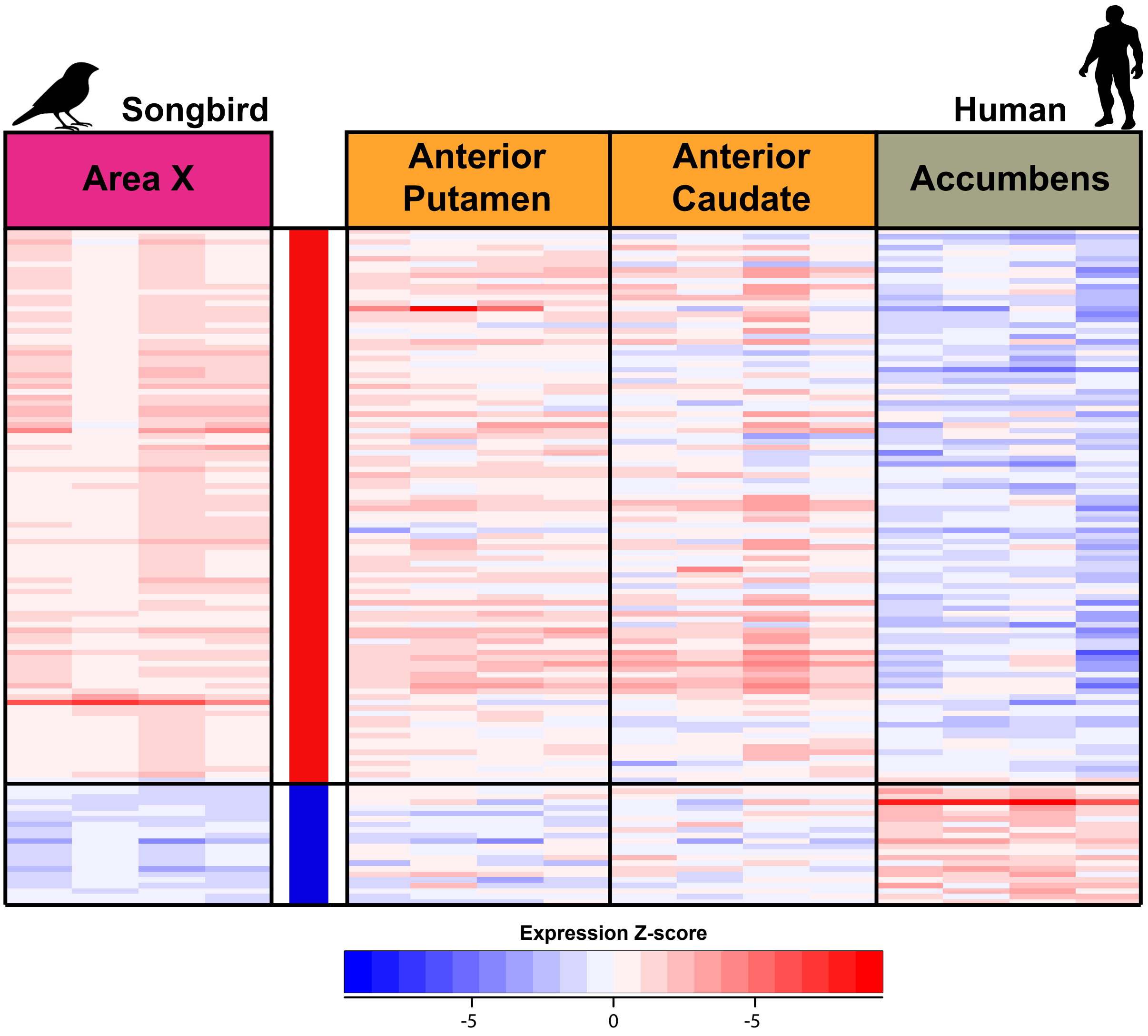


**Figure S9: Heatmap visualizing z-score normalized expression of the convergent gene set in songbird Area X (left) and human anterior striatum (right).** Each column is a replicate of the above brain region. Each row is a gene exhibiting convergent expression across songbird and human vocal motor striatal regions. Top boxes group and color-code each brain region. The three human striatal regions (putamen, caudate, accumbens) are shown separately. A total of 118 genes exhibit convergent expression between songbird Area X and human putamen.

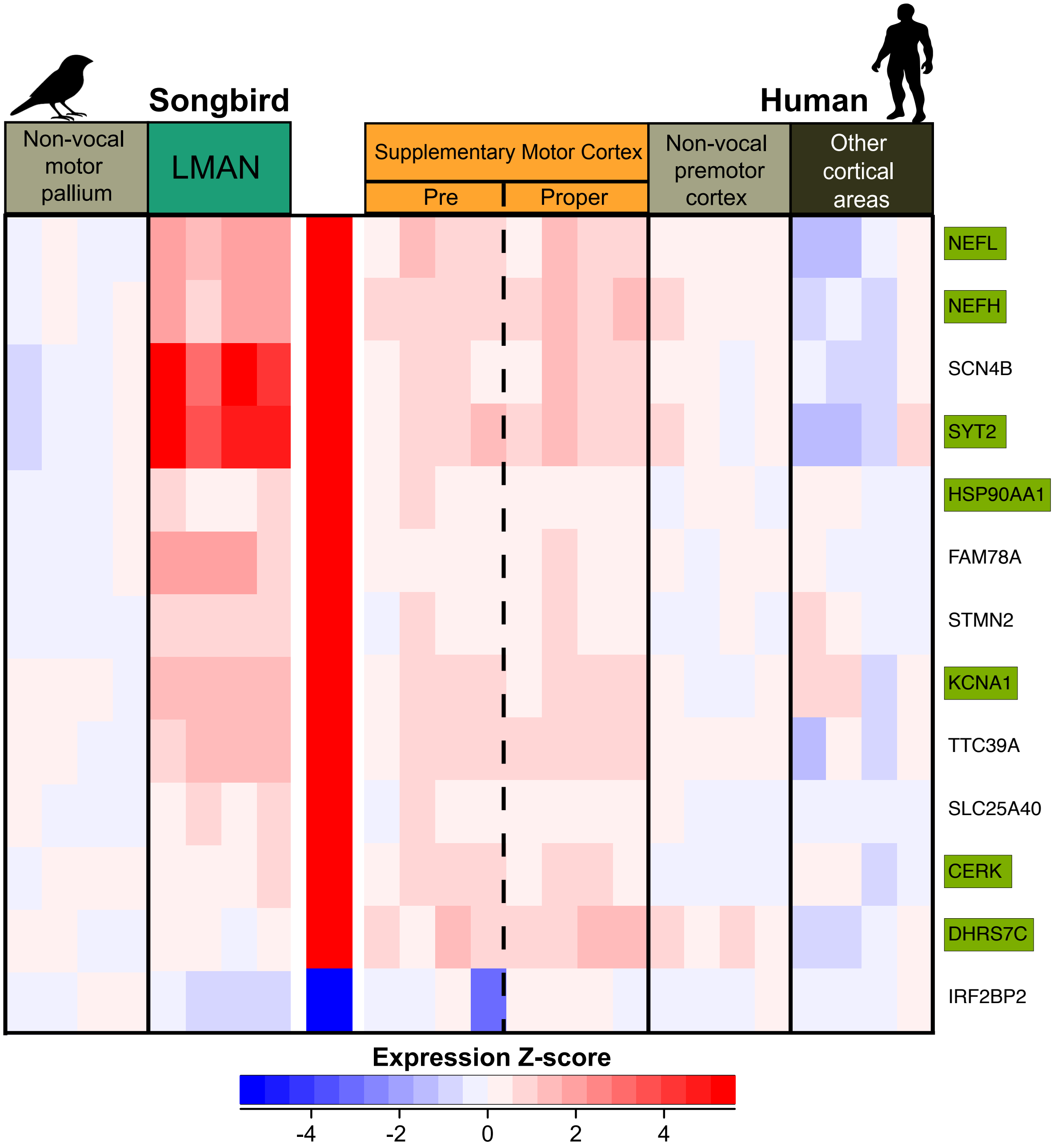


**Figure S10: Convergent molecular specializations between songbird HVC/LMAN and human SMA.** Heatmap visualizing z-score normalized expression of the convergent gene set in songbird LMAN (left) and human SMA (right). Each column is a replicate of the above brain region. Each row is a gene exhibiting convergent expression across songbird and human vocal motor regions. Non-vocal motor regions and other cortical regions (human) for each species are included as controls. The two subregions of the SMA are also delineated (dotted lines). All genes (n=12) exhibit convergent expression between songbird LMAN and human SMA, with nearly all exhibiting upregulation. Genes exhibiting convergence with HVC and SMA (green boxes) were all similarly specialized in LMAN.

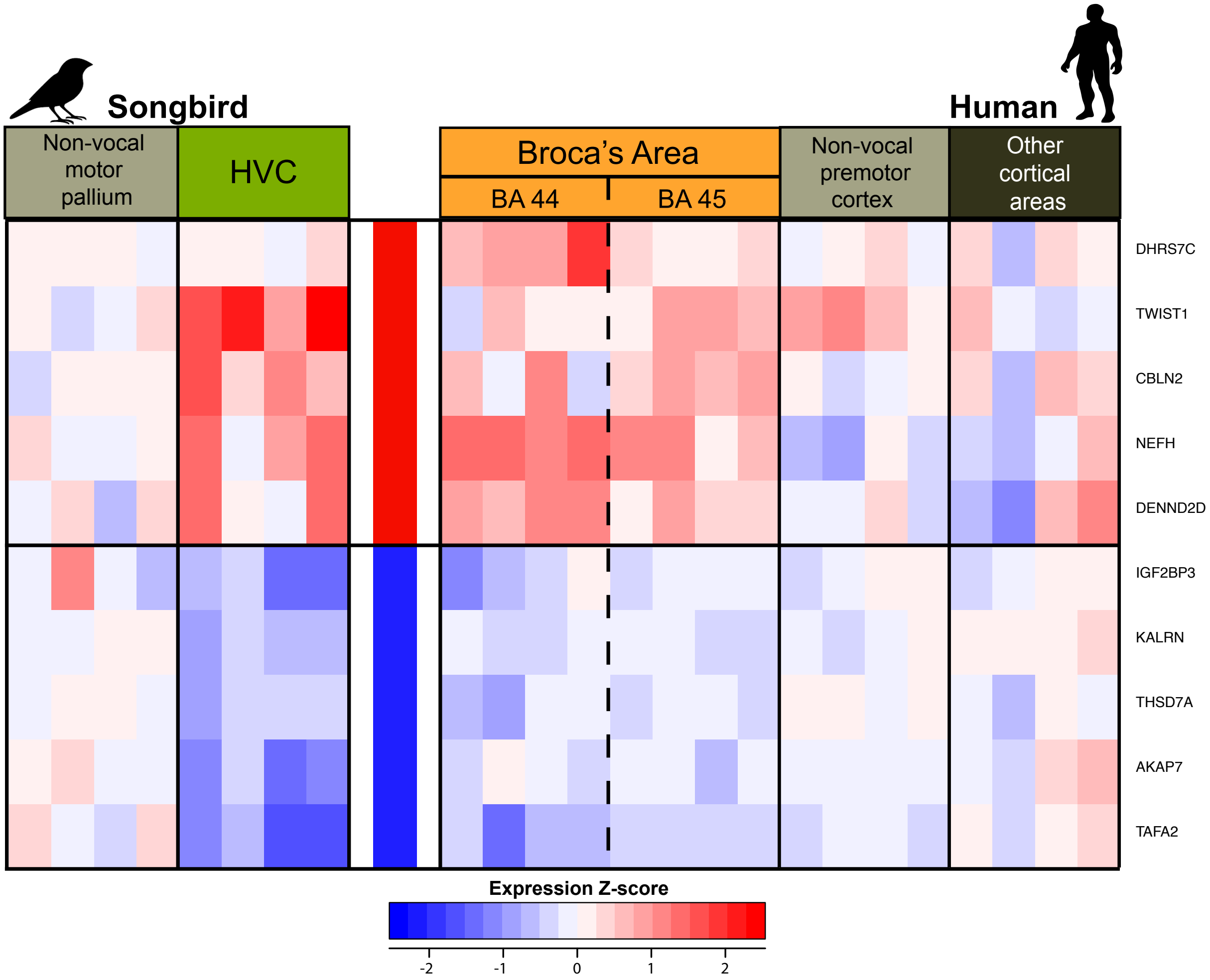


**Figure S11: Convergent molecular specializations between songbird HVC and human Broca's area.** Heatmap visualizing z-score normalized expression of the convergent gene set in songbird HVC (left) and human Broca's area (right). Each column is a replicate of the above brain region. Each row is a gene exhibiting convergent expression across songbird and human vocal motor regions. Non-vocal motor regions and other cortical regions (human) for each species are included as controls. The two subregions of the Broca's area are also delineated (dotted lines). These genes (n=10) exhibit convergent expression between songbird HVC and human Broca's, with a 50/50 split of up/down regulation. GO analysis revealed no significant functional enrichment for this gene set.

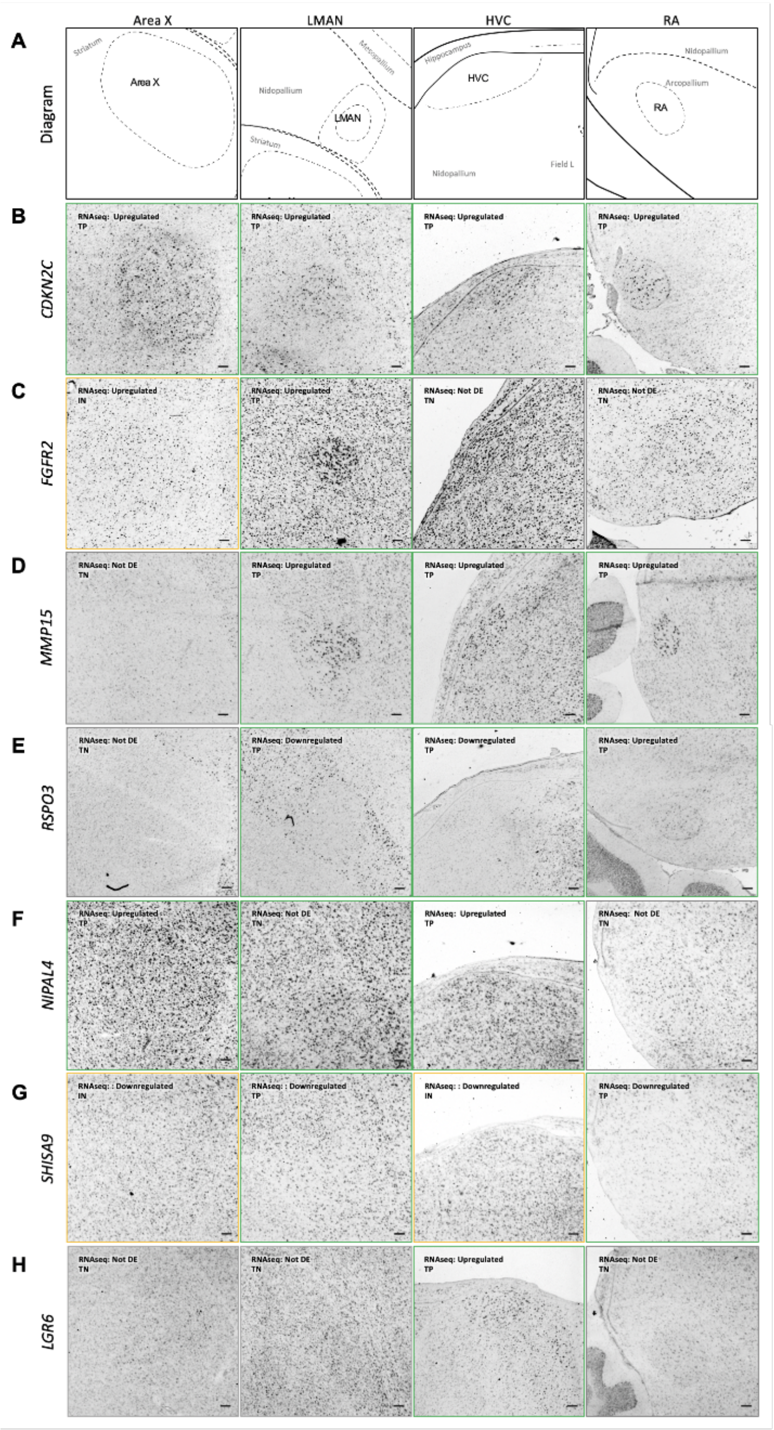


**Figure S12: Validation of convergent DEGs between humans and songbirds**. (**A**) Approximate anatomical diagram of adult male zebra finch vocal learning song nuclei. (**B-H**) Colorimetric *in situ* hybridization characterizing (**B**) *CDKN2C*, (**C**) *FGFR2*, (**D**) *MMP15*, (**E**) *RSPO3*, (**F**) *NIPAL4*, (**G**) *SHISA9*, and (**H**) *LGR6*. Image outlines are color-coded according to comparison with RNA-seq data. True Positive (TP) = Green; True Negative (TN) = Grey; False Positive (FP) = Red; False Negative (FN) = Blue; Inconclusive (IN) = Yellow.

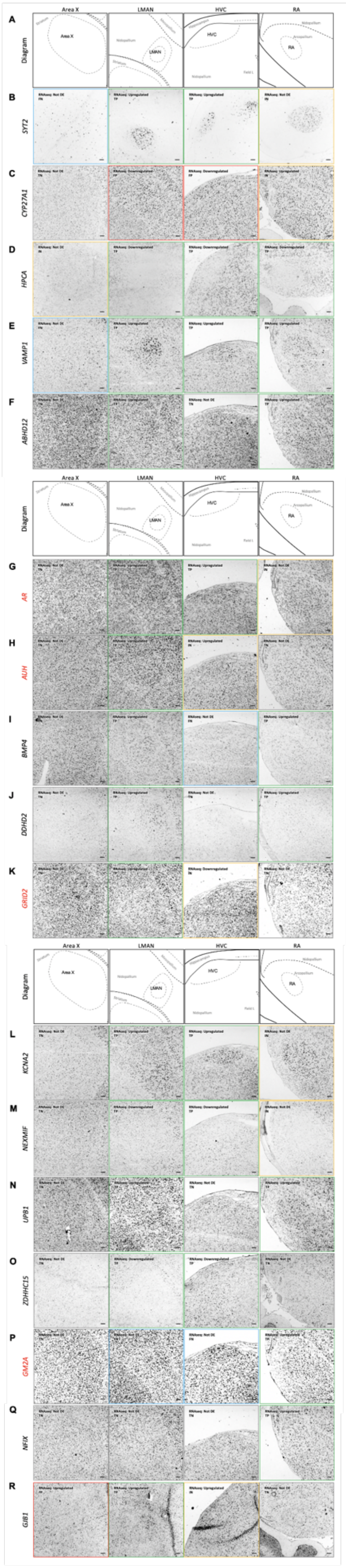


**Figure S13: Validation of song nuclei DEGs convergent with human LMC DEGs, with relevance to human speech disorders.** (**A**) Approximate anatomical diagram of adult male zebra finch vocal learning song nuclei. Colorimetric *in situ* hybridization characterizing (**B**) *SYT2*, (**C**) *CYP27A1*, (**D**) *HPCA*, (**E**) *VAMP1*, (**F**) *ABHD12*, (**G**) *AR*, (**H**) *AUH*, (**I**) *BMP4*, (**J**) *DDHD2*, (**K**) *GRID2*, (**L**) *KCNA2*, (**M**) *NEXMIF*, (**N**) *UPB1*, (**O**) *ZDHHC15*, (**P**) *GM2A*, (**Q**) *NFIX*, and (**R**) *GJB1*. Image outlines are color-coded according to comparison with RNA-seq data. True Positive (TP) = Green; True Negative (TN) = Grey; False Positive (FP) = Red; False Negative (FN) = Blue; Inconclusive (IN) = Yellow.

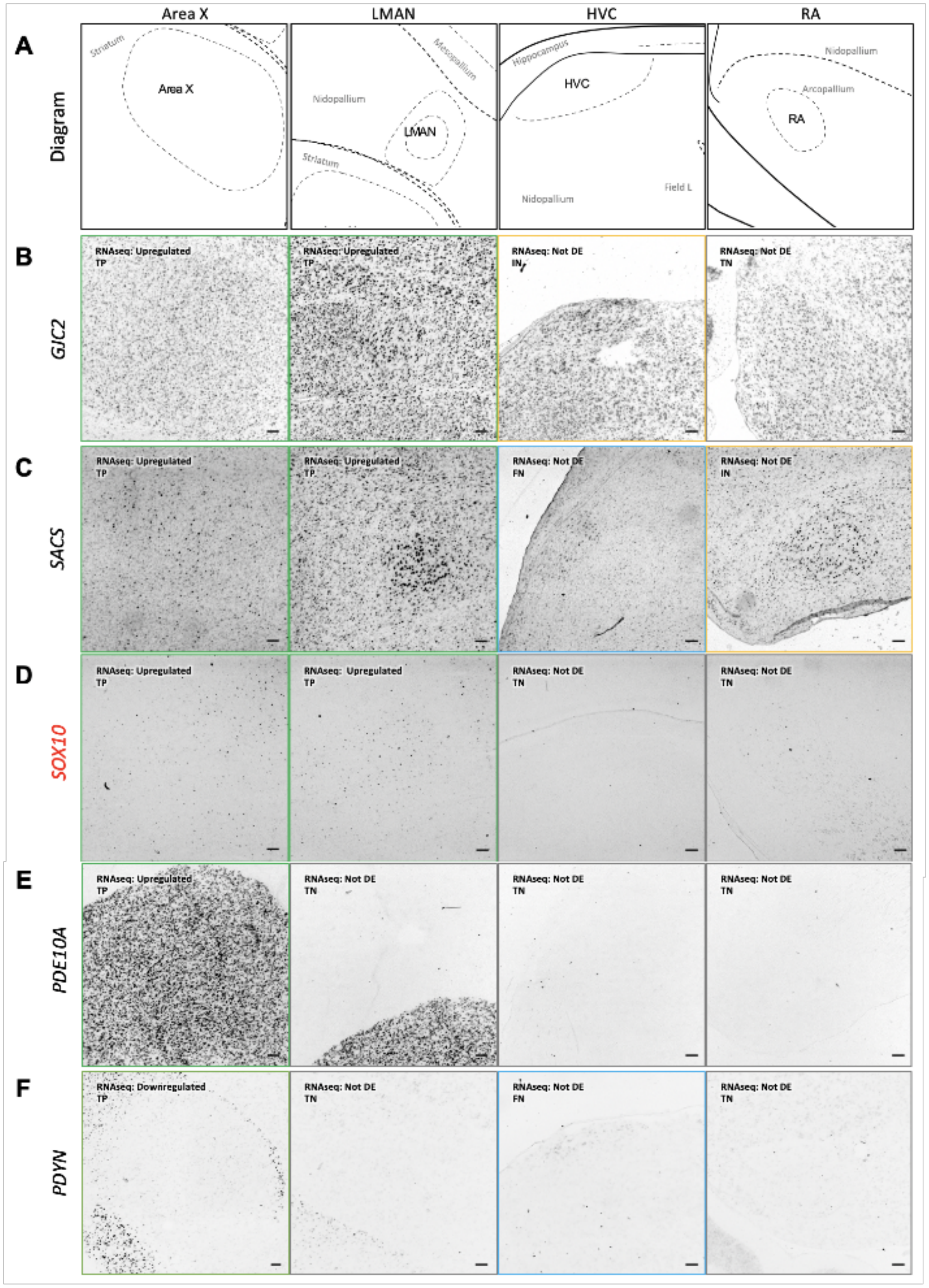


**Figure S14: Validation of song nuclei DEGs convergent with human Striatum DEGs, with relevance to human speech disorders.** (**A**) Approximate anatomical diagram of adult male zebra finch vocal learning song nuclei. Colorimetric *in situ* hybridization of, characterizing, (**B**) *GJC2*, (**C**) *SACS*, (**D**) *SOX10*, (**E**) *PDE10A*, and (**F**) *PDYN*. Image outlines are color-coded according to comparison with RNA-seq data. True Positive (TP) = Green; True Negative (TN) = Grey; False Positive (FP) = Red; False Negative (FN) = Blue; Inconclusive (IN) = Yellow.

**Table S1: Differential gene expression in the zebra finch song system.** (**1**) Summary of differential gene expression analysis results (FDR < 0.5). A "1" denotes upregulation, a "-1" denotes downregulation, and a "0" denotes no change relative to the adjacent non-vocal brain region to the given nucleus. More detailed results for Area X (**2**), LMAN (**3**), HVC (**4**), and RA (**5**) are provided, including log2 fold changes and adjusted p values.

**Table S2: Validation of differential expression with *in situ hybridization* in zebra finch brains.** (**1**) For each gene (n = 178), the differential expression results for each nucleus were summarized as in **Table S1**, as well as the call determined by the blinded evaluators if the gene was a true positive (TP), false positive (FP), true negative (TN), false negative (FN), or inconclusive (IN). (**2**) Summary of in situ validation rates for each nucleus.

**Table S3: Allen Human Brain Atlas microarray metadata**. The data table consists of 3,681 unique samples (**1**) from six individuals each measuring 18,416 genes (**2**). All sample coordinates are provided in MNI space, along with chromosome location and Ensembl ID for each gene.

**Table S4: Human speech region and background sample metadata.** The data table consists of 1,432 cortical (**1**) and 123 striatal (**2**) samples from **Table S3** annotated by the brain region in which they reside which is active during human speech/language tasks. All samples labeled as "non" denote regions not activated by speech and were used as background in the given comparison.

**Table S5: Differential gene expression in the human vocal motor system. (1-17)** Results are named for each human brain region examined in this study (**Fig. 3A-D)** relative to either the cortex (1-13) or striatum (14-17). Differential expression (DE) status is coded as described previously.

**Table S6: Gene sets exhibiting convergent expression between songbird and human brain regions. (1-14)** Lists of genes showing convergent expression between human and songbird brain regions. Ensembl IDs from both species as well as direction of specialized expression are reported.

**Table S7: Gene ontology analysis of convergent gene lists in songbird and human. (1-10)** Results from gene ontology analysis conducted with gprofileR2 using the list of one-to-one songbird and human orthologs as a custom background. Only significant results (FDR < 0.05) are reported from the 3 major GO categories, biological process (BP), cellular compartment (CC), and molecular function (MF). Any gene list from **Table S6** that doesn't appear here was not significantly enriched for any known function.

**References and Notes: Supplementary Materials**


69. G. Gedman *et al.*, As above, so below: Whole transcriptome profiling demonstrates strong molecular similarities between avian dorsal and ventral pallial subdivisions. *J Comp Neurol*, (2021).
70. G. Feenders *et al.*, Molecular mapping of movement-associated areas in the avian brain: a motor theory for vocal learning origin. *PLoS ONE* **3**, e1768 (2008).
71. A. Rhie *et al.*, Towards complete and error-free genome assemblies of all vertebrate species. *Nature* **592**, 737-746 (2021).
72. A. R. Pfenning *et al.*, Convergent transcriptional specializations in the brains of humans and song-learning birds. *Science* **346**, 1256846 (2014).
73. M. Belyk, P. Q. Pfordresher, M. Liotti, S. Brown, The Neural Basis of Vocal Pitch Imitation in Humans. *J Cogn Neurosci* **28**, 621-635 (2016).
74. S. Köhler *et al.*, The Human Phenotype Ontology in 2021. *Nucleic Acids Research* **49**, D1207-D1217 (2020).
75. P. V. Lovell *et al.*, ZEBrA: Zebra finch Expression Brain Atlas-A resource for comparative molecular neuroanatomy and brain evolution studies. *J Comp Neurol* **528**, 2099-2131 (2020).
76. M. T. Biegler, L. J. Cantin, D. L. Scarano, E. D. Jarvis, Controlling for activity-dependent genes and behavioral states is critical for determining brain relationships within and across species. *J Comp Neurol*, (2021).
77. T. E. Bakken *et al.*, Evolution of cellular diversity in primary motor cortex of human, marmoset monkey, and mouse. *bioRxiv*, 2020.2003.2031.016972 (2020).
78. B. M. Colquitt, D. P. Merullo, G. Konopka, T. F. Roberts, M. S. Brainard, Cellular transcriptomics reveals evolutionary identities of songbird vocal circuits. *Science* **371**, eabd9704 (2021).
79. W. C. Warren *et al.*, The genome of a songbird. *Nature* **464**, 757-762 (2010).
80. J. Korlach *et al.*, De novo PacBio long-read and phased avian genome assemblies correct and add to reference genes generated with intermediate and short reads. *Gigascience* **6**, 1-16 (2017).
81. M. J. Hawrylycz *et al.*, An anatomically comprehensive atlas of the adult human brain transcriptome. *Nature* **489**, 391-399 (2012).
82. J. W. Bohland, F. H. Guenther, An fMRI investigation of syllable sequence production. *Neuroimage* **32**, 821-841 (2006).
83. A. J. Simmonds, R. Leech, P. Iverson, R. J. Wise, The response of the anterior striatum during adult human vocal learning. *J Neurophysiol* **112**, 792-801 (2014).
84. R. Báez-Mendoza, W. Schultz, The role of the striatum in social behavior. *Front Neurosci* **7**, 233 (2013).
85. K. Simonyan, U. Jürgens, Efferent subcortical projections of the laryngeal motorcortex in the rhesus monkey. *Brain Res* **974**, 43-59 (2003).
86. S. Lehéricy *et al.*, Diffusion tensor fiber tracking shows distinct corticostriatal circuits in humans. *Ann Neurol* **55**, 522-529 (2004).
87. S. Kumar *et al.*, A Brain System for Auditory Working Memory. *J Neurosci* **36**, 4492-4505 (2016).
88. M. Inase, J. A. Buford, M. E. Anderson, Changes in the control of arm position, movement, and thalamic discharge during local inactivation in the globus pallidus of the monkey. *J Neurophysiol* **75**, 1087-1104 (1996).

89. U. Jurgens, Neural pathways underlying vocal control. *Neurosci Biobehav Rev* **26**, 235-258 (2002).
90. E. D. Jarvis, Evolution of vocal learning and spoken language. *Science* **366**, 50-54 (2019).
91. L. Kubikova *et al.*, Basal ganglia function, stuttering, sequencing, and repair in adult songbirds. *Sci Rep* **4**, 6590 (2014).
92. C. S. Lai, S. E. Fisher, J. A. Hurst, F. Vargha-Khadem, A. P. Monaco, A forkhead-domain gene is mutated in a severe speech and language disorder. *Nature* **413**, 519-523 (2001).
93. I. Teramitsu, S. A. White, FoxP2 regulation during undirected singing in adult songbirds. *J Neurosci* **26**, 7390-7394 (2006).
94. S. Haesler *et al.*, Incomplete and inaccurate vocal imitation after knockdown of FoxP2 in songbird basal ganglia nucleus Area X. *PLoS Biol* **5**, e321 (2007).
95. I. Hertrich, S. Dietrich, H. Ackermann, The role of the supplementary motor area for speech and language processing. *Neurosci Biobehav Rev* **68**, 602-610 (2016).
96. P. A. Chouinard, T. Paus, What have We Learned from "Perturbing" the Human Cortical Motor System with Transcranial Magnetic Stimulation? *Front Hum Neurosci* **4**, 173 (2010).
97. A. Alm, *Cluttering: a neurological perspective*. (Psychology Press, ed. 1, 2011).
98. P. Broca, Nouvelle observation d'aphemie produite par une lesion de la moitie posterierure des deuxieme et troisieme circonvolutions frontales. *Bulletin de la Societe Anatomy de Paris* **VI**, 398-407 (1861).
99. S. Bookheimer, Functional MRI of language: new approaches to understanding the cortical organization of semantic processing. *Annu Rev Neurosci* **25**, 151-188 (2002).
100. G. A. Castellucci, C. K. Kovach, M. A. Howard, J. D. W. Greenlee, M. A. Long, A speech planning network for interactive language use. *Nature* **602**, 117-122 (2022).
